# The study of differential expressions of MCPH and Seckel syndrome genes and their paralogues

**DOI:** 10.1101/2025.05.21.655339

**Authors:** Muhammad Mohsin, Sana Riaz, Mohammad Imran, James J Cox, Maryam Khurshid

## Abstract

Seckel syndrome and MCPH are single-gene neurodevelopmental disorders, with or without dwarfism. These genes are parts of essential cellular pathways but are regulated differently in both forms of microcephaly. These genes have diverged from single ancestral genes in to related but different gene paralogues during evolution. It is suggested that the paralogues might be redundant in other body organs and might rescue any abnormality in cases of MCPH, but not in Seckel syndrome. This study uncovers the spatio-temporal dynamics of these genes together with their paralogues at the mRNA level where the different tissues of embryos might exhibit different transcript variants or different levels of the various transcript variants to find the complementation in case of the MCPH but divergence in case of the Seckel syndrome. Here, we studied Cdk5Rap2-Pde4dip, Phc1-Phc3 (for MCPH) and Cep63-Ccdc67 (for Seckel) in the embryonic mouse tissues using RT-PCR throughout the peak of neurogenesis from E12.5 to E18.5. We found already known and novel spatio-temporal differential expression suggesting different regulation at the mRNA level. The results of PCR were analyzed on the agarose gel and were quantified by assigning relative intensity percentage scores to each band and plotting the cumulative expression of each gene in to the graphs for spatio-temporal dynamics and plotted each gene separately for temporal dynamics. For MCPH, Cdk5Rap2-Pde4dip had similar profiles and seemed to be redundant whereas Phc1-Phc3 seem to have complementary roles in brain development. The expression profiles of Cep63-Ccdc67 were very different indicating of their divergent and essential roles in development.

## Introduction

Prenatal and postnatal microcephaly has been reported in humans. This might be due to genetic factors or genes that might cause non-syndromic microcephaly, (Thornton and Woods, 2009a) where only head size is reduced or syndromic forms where all organs are small together with craniofacial and other abnormalities(Asif *et al*., 2023a; Khetarpal *et al*., 2016). Autosomal recessive primary microcephaly (MCPH) is a non-syndromic condition in which the affected have brain size is reduced together with mild to moderate mental retardation, and the rest of the organs in the body are of normal size(Zaqout and Kaindl, 2022a). Seckel syndrome on the other hand, is syndromic microcephaly which occurs in utero but is progressive and shows severe microcephaly with dwarfism and other anomalies (Khetarpal *et al*., 2016; Nano and Basto, 2017). Both MCPH and Seckel syndrome are heterogeneous and hence, indicate the different mechanisms of pathogenesis resulting from the gene mutations that may which sometimes overlap such as MCPH6 gene CENPJ, which is also involved in the Seckel syndrome(Asif *et al*., 2023a; Cuccurullo *et al*., 2022). Both are heterogeneous, and single genes are causative agents of both, the agents that are inherited in an autosomal recessive manner, so there must be some differential regulation done by the genes in each category and must control development in different manners(Asif *et al*., 2023a; Kaindl *et al*., 2010a). These genes are targeted to several core pathways such as centriole biogenesis, centrosome/spindle pole, kinetochore-microtubular pathway, nuclear pathways including the condensation-decondensation and nuclear envelope dynamics and other sub-cellular pathways(Bakircioglu *et al*., 2011; Nicholas *et al*., 2010a; Sir *et al*., 2011a; Thornton and Woods, 2009a).

During evolution genes diverge from ancestral genes in to different molecular entities with different acquired functions that may be related as these retain a considerable similarity termed as gene paralogues(Castellanos *et al*., 2024). Primary microcephaly genes have been studied in the past decade for expression and by gene knock out in animal models which have provided a large insight in to the disease but the data suggests ubiquitous roles of these proteins in all body cells and mouse knockout are not true models of the disease with small to no reduction of organs(Perlman, 2016). Furthermore, the comparative expression analysis of genes and their paralogues might answer the question. Some studies have been performed with a double knockout of more than one PM gene and found a severe phenotype indicative of involvement in one master pathway of the cell incorporating all dynamics such as the cell cycle, centrosomal/spindle pole, centriole biogenesis, nucleus and others where these genes have established roles(Farcy *et al*., 2023; Jayaraman *et al*., 2016a; Morris-Rosendahl and Kaindl, 2015).

One aspect that was missing was the differential expression patterns of microcephaly genes and their paralogues, which can be explored for spatio-temporal expression analysis together with their established paralogues in both forms of microcephaly(Duan *et al*., 2022a; Hettige and Ernst, 2019). Here we selected *CDK5RAP2-PDE4DIP* and *PHC1-PHC3* genes and their respective paralogues duos for MCPH and *CEP63-CCDC67* genes from Seckel syndrome(Awad *et al*., 2013a; Kraemer *et al*., 2015; Ma *et al*., 2014; Mani, 2022; Sir *et al*., 2011a; Tonkin *et al*., 2002a; Zhao *et al*., 2013). These genes were selected because of their targeting involves different pathways that might be differentially important at different embryonic days. We used Swiss albino mice for the study and performed mating via a timed mating protocol and to obtain staged embryos(Byers *et al*., 2012a). Later, we employed RT-PCR using gene specific primers to determine if there are some variations in transcripts and/or the level of expression of the transcripts.

*CDK5RAP2* at MCPH5 is involved in centrosome dynamics whereas *PHC1* at MCP11 is involved in nuclear dynamics, while the Seckel gene, *CEP63* at SCKL6, is a core centrioler gene and is involved in centriole duplication, and their respective paralogues have related functions in the same or different pathways (Zaqout and Kaindl, 2022a). In order to see if genes and their paralogues complement each other or if they replace each other’s function being redundant or/and completely diverged and have different roles we set out to understand via simple but informative RT-PCR analysis an established method to uncover differential transcript variants or/and levels of expression.

## Materials and Methods

### Animal mating and tissue harvesting

Swiss albino mice strain (BALB/cJ) was used and the mice were purchased from the Department of Pharmacy at Islamia University of Bahawalpur. These mice were of wildtype and were inbred. Housing was performed according to international standards. Timed mating of mice (Imran *et al*., 2024) was done at proestrous-estrous stage and staged embryos were collected later at specific required days in RNA later^TM^ (Cat. No. AM7020, Invitrogen by Thermo Fisher Scientific). The acquisition of embryos required dissection of the pregnant mice and it was done by intraperitoneal injection of dissociative agent, ketamine in combination with α_2_-adrenergic receptor agonist, xylazine according to the weight of the pregnant mice (ketamine 100mg/kg and xylazine 10mg/kg of mice weight). The xylazine-ketamine was injected in a 1:10 ratio combination to prevent pain sensing, to induce sedation and to act as anesthetic and muscle relaxant. The mice were left for half an hour and checked physically for vital signs such as heart beat and sensation and ensured that they are expired before proceeding. The embryonic tissues are sensitive and to ensure that tissues are not contaminated with chemical agents, we performed cervical dislocation and within this ensured the death of animal as well as no residual pain sensing. The study was approved by institutional ethical committee of Islamia University of Bahawalpur (letter no. 3990/ORIC dated 26/04/2021).

Two group of mice, timed mating group and blind mating group, were compared and this was done to investigate the efficiency of the pregnancies in small groups to acquire staged embryos for the project. Ten cages, 4 for female mice which were housed together to synchronize their estrous cycle and separate cages were allotted for all the male mice. Ten female mice and 5 male mice in each group, which makes total 15+15=30 mice in the experiment. It was decided by the available space in the animal housing facility which is not very spacious and it is the only one facility available on campus. The healthy virgin females were preferred whereas, stud male mice were inducted. The mice were not undergoing any drug trial and were only for this project. The male mice were housed in separate cages but females were housed in groups of 3-4 to synchronize their estrous cycle. The mice were housed in wooden cages with sawdust bedding and water bottles and were placed in regulated temperature and light. The male mice fight if kept together whereas the female mice can be kept together. So, we kept male mice separate and females were shifted in their cages for mating as planned in the experiment. The mice were mated in female to male mice ratios of 2:1. The mating was done in dark under cooler conditions. The appearance of vaginal plug next day early in the morning was the indication to separate the females and after counting the embryonic days they were sacrificed at required day. The day of the plug is E0.5. The first group random/blind mating was used to recover the embryos after mating and is not dependent on the day of estrous cycle in female mice or any other regulation before mating. In random/blind mating group any day for setting up the mating was fine. In timed mating group female mice estrous cycle was tracked by vaginal cytology and visual observation and were left to mate when mice were at pro-estrous/estrous cycle stage. This was done to compare the yield of staged embryos in each group. The number of pregnancies were more in timed mating than random mating. We recovered total of 4 pregnant mice in blind mating trail and 8 in timed mating trial. The null hypothesis was investigated and was proven correct and sample size was calculated that was required minimally in small laboratory settings. Two proportion Z test was used to determine any statistical difference between two mating strategies. The *p* value was 0.06 which is not significant. This calculation was done on the Microsoft Excel sheet. It does not meet the assumption as previous studies found the timed mating strategy to be statistically significant for pregnancies that random mating (Byers *et al*., 2012b; Dorsch *et al*., 2020). However, our number of mice was not very large. In total, at E7.5 =1, E12.5 = 1, E13.5 =3, E14.5 =3, E15.5 =3 and E18.5 = 1 were recovered (More details on the complete experiment are available at (Imran *et al*., 2024)).

The embryos collected in the study were used in this study and we did mRNA expression profiling. From each pregnant mice > 6 embryos were recovered which were separated in to head regions, Hd, Rest of embryos, RE (embryonic body) and some were kept whole as whole embryos, WE, all were collected in RNA Later^TM^ (Cat. No. AM7020, Invitrogen by Thermo Fisher Scientific). E7.5 was too early for neurogenesis and was not included in the analysis. Peak of neurogenesis is from E12.5 to E15.5 and late stage of neurogenesis is at E18.5 and neurodevelopment disorders of MCPH and Seckel syndrome are best studied at these stages to capture spatio-temporal dynamics. The staging was confirmed via Immunohistochemistry of paraffin embedded embryos of these stages by use of molecular markers. For this paper, we did three replicates for each stage (where-ever possible) and tissues (Hd, RE and WE) however, some samples were missing due to loss of tissue samples during optimization.

### RNA Extraction

Total RNA was extracted from 100mg of tissue using the TRIzol (Cat. No. 15596018, Ambion by Life technologies) method. The minced tissue was incubated with 1mL of TRIzol reagent overnight at 37°C. The next day, the tubes were centrifuged at 10,000 RPM for 20 mins at 4°C. The supernatant was separated in to the aqueous phase containing RNA and organic phase containing DNA and Proteins by using chloroform (GPR^®^). The aqueous phase was taken and RNA was pelleted down by isopropanol (Sigma Aldrich) incubation and centrifugation at 10,000 RPM for 20 mins. The pellet was washed with 75% Ethanol (Sigma Aldrich) and the samples were centrifuged for 5 mins at 7000 RPM at 4°C, supernatant was discarded and pellets were air dried at room temperature for 5 mins. The pellet was resuspended in 20µl of RNase/DNase free water in 65°C incubator for 5-10 mins. RNA was stored at −20°C.

### First strand cDNA synthesis

The nanodrop method (Gen5) was used to calculate absorbance at 260nm and 280 nm, 260/280 ratios were used to select good quality of RNA in terms of purity and integrity (Gen5 software). The values were used to calculate 1µg RNA in each sample and first strand cDNA was synthesized by RevertAid (Cat. No. EP0442, Thermo Fisher Scientific) first-strand cDNA synthesis kit. The reverse transcription was done using the described protocol provided with the kit. 18mer, Oligo dT primers supplied in the kit were used to amplify the first strand. Blank and RT-ve controls were used.

### Second strand cDNA synthesis

The synthesized first-strand cDNA synthesized was amplified into double stranded DNA by using gene specific primers and the estimated known sizes were up to 900bp so Taq DNA polymerase system (Cat. No. EP0402 by Thermo Fisher Scientific or Cat No. MB101-0500 by GeneDirex) was used as a standard method.

### Gene specific primers

Intron-spanning gene-specific primers targeting all known common transcript variants and sometimes specific transcripts were used. Primer 3plus and primer blast were used for the primer designing. The Insilico-PCR was done to calculate the product sizes and targeted transcripts from the UCSC genome browser. The melting temperatures and GC content were also calculated. The sequences are shown in the supplementary tables (Supplementary Table, S2). The primers were synthesized by oligo synthesizing company, the Macrogen, South Korea upon ordering. The lyophilized primers were shipped at room temperature and were resuspended in DNase/RNase free water to make 100mM stocks. In PCR, 0.1 to 0.2μM concentrations were used for primer pairs whichever was suitable. Blank was used with all primers and adult brain as positive control was used. The Gapdh control primer was used and together with samples, Gapdh positive RNA control was also done.

### PCR program

PCR was performed in the gradient thermal cycler (Ref. A37029, Applied Biosystems by Thermo Fisher Scientific). PCR was done at initial denaturation at 94°C for 5 mins, 35 cycles of denaturation at 94°C for 30 sec, annealing at 53/55/56°C for 30 sec and extension at 72°C for 1 min. The final extension step was done at 72°C for 10 mins. The PCR machine was set at 4°C forever at the end of program.

### Gel electrophoresis

A 2% agarose (R0492 by Thermo Fisher Scientific) gel (90mL) was prepared and 3µl ethidium bromide was added for visualization in a UV transilluminator (UV Tech). A 100 bp DNA ladder (FM0323 by Thermo Fisher Scientific) was used with each primer to determine the size of the bands. The gel was run in horizontal gel tank (Ref. MU-E73-91338 by Stalwart) at 50V for 30-40 mins at RT in 1XTBE buffer.

### Relative Intensity scoring

DNA bands were marked by their presence and relative intensity. The highest intensity was scored as 4 (100%) and was further calculated down to 1(25%) as the intensity in the samples drop. The absence of bands was numbered 0 (0%). Later, the cumulative scores of all bands for each primer were added in each sample, and relative percentages were calculated for each sample (Supplementary Table, S1). Means were calculated for more than one sample wherever applicable.

### Excel software

The Microsoft Excel was used to plot the relative percentages of each sample in to bar graphs and individual line graphs. Error bars were also added, where required.

## Results

### CDK5RAP2 and PDE4DIP

CDK5RAP2 at MCPH3(Zaqout and Kaindl, 2022a) is one of the first few MCPH genes to be identified. CDK5RAP2 and PDE4DIP/Myomegalin (Mani, 2022) were identified and confirmed as paralogues of each other by Position Specific Iteration BLAST, PSI-BLAST program (see figure no. 1, A-C). The multiple sequence alignment, MSA showed very high similarities between the two proteins throughout most regions of the two proteins (see figure no. 1, A). These were further subjected to spatio-temporal investigation (Duan *et al*., 2022a; Vaid and Huttner, 2020a) in mice staged embryos in order to determine whether they can actually act by replacing each other. Cdk5Rap2, a protein kinase involved in the centrosome dynamics, was found to be most highly expressed in the head of E14.5 mice, and interestingly, its paralogue PDE4DIP was also most high in the E14.5 mice (see figure no. 2, A-D and 3, A-D). At E14.5, the basal progenitors are abundant and it is the mid-neurogenesis peak indicating that both genes are essential most specifically to the basal progenitors. Furthermore, we observed the expected transcript variants in the samples, as well as some unexpected bands for potential new transcript variants. We plotted the cumulative expression as well as only the expected expression of transcript variants by intensity scoring, and also the potential new transcript variants were treated in the same way. All of these different scenarios gave us a similar profile where the expression of *Cdk5Rap2* gradually up-regulated after E12.5 and E13.5, and became highest in the E14.5 head, and stayed at the same level at E15.5 and E18.5, in the mouse embryonic heads suggesting a very important role of Cdk5Rap2 in these stages when basal progenitors are most abundant and SVZ is the major layer in the neuroepithelium (see figure no. 2 B-D). Its paralogue, *PDE4DIP*, showed the highest expression of all the transcript variants both expected and novel, in the E13.5, followed by E14.5. However, in the potentially new variants, E14.5 was the highest and the expression was downregulated in all scenarios later in neurogenesis (see figure no. 3D). This is very interesting, as at E13.5 and E14.5, there are apical as well as basal progenitors, with the latter being abundant, and it might be possible that the embryonic development in mice is dependent on the expression of genes and there paralogues in the brain, where both are required for brain development and are complementary, but the time window for the paralogue expression is short compared with the MCPH gene, *CDK5RAP2* suggesting that the prolonged role of this gene is required for normal brain development, which is also the longest stage in the human development.

**Figure 1:**
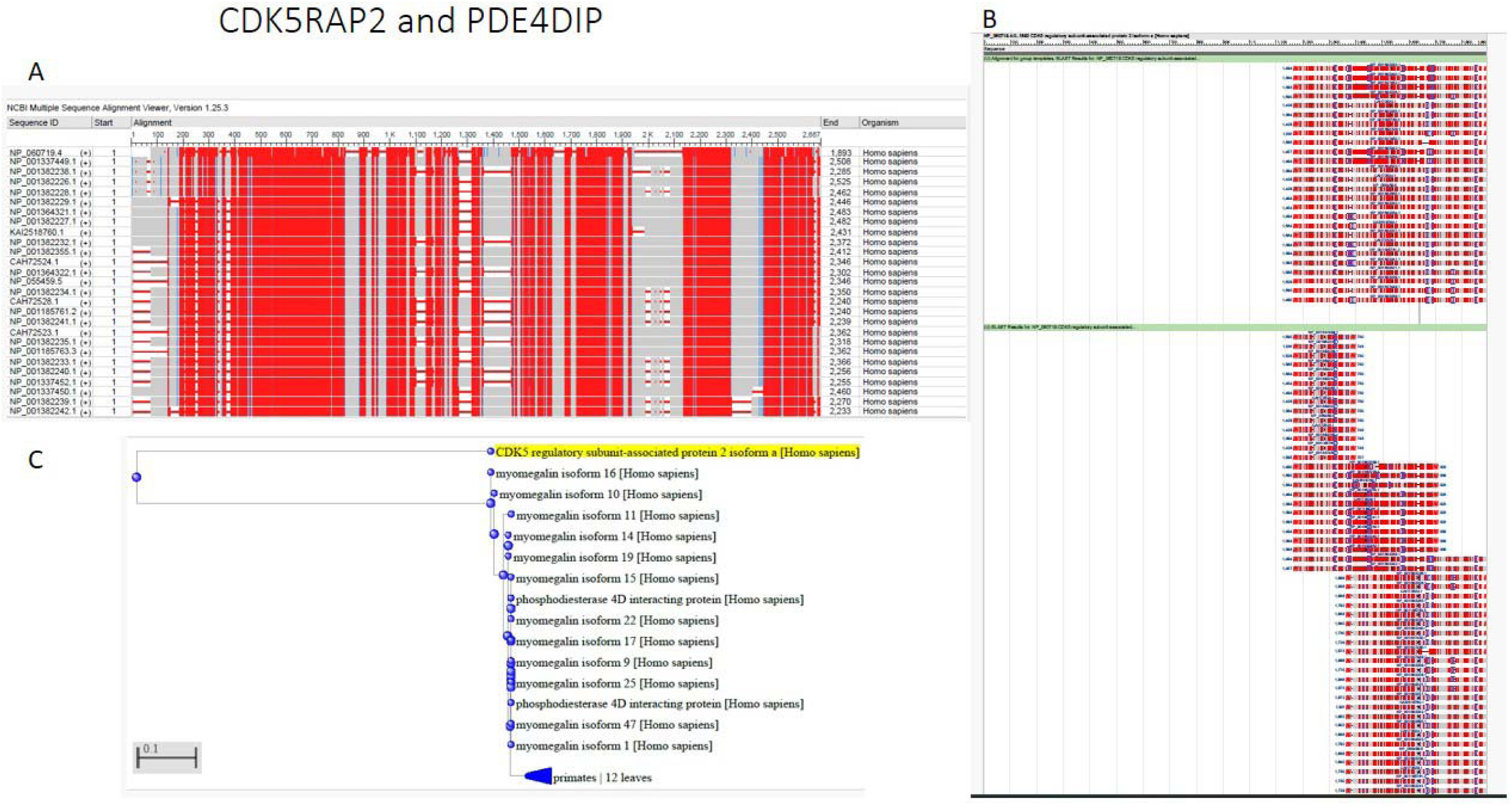
PSI BLAST was run to identify and confirm the ancestry, similarity and overall similarity between CDK5RAP2 and PDE4DIP/Myomegalin in Humans. A) Multiple Sequence Alignment between the CDK5RAP2 and Myomegalin/PDE4DIP is shown and is indicative of very high similarity especially at the N-terminus and other domains, B) BLAST between Isoform 1 of CDK5RAP2 was done and came up with similarities in various regions with almost all isoforms of Myomegalin/PDE4DIP, C) Ancestry of both CDK5RAP2 and PDE4DIP was confirmed by Gene tree where both genes split in a single evolution event and myomegalin showed a lot of further splits indicative of active evolution.

**Figure 2:**
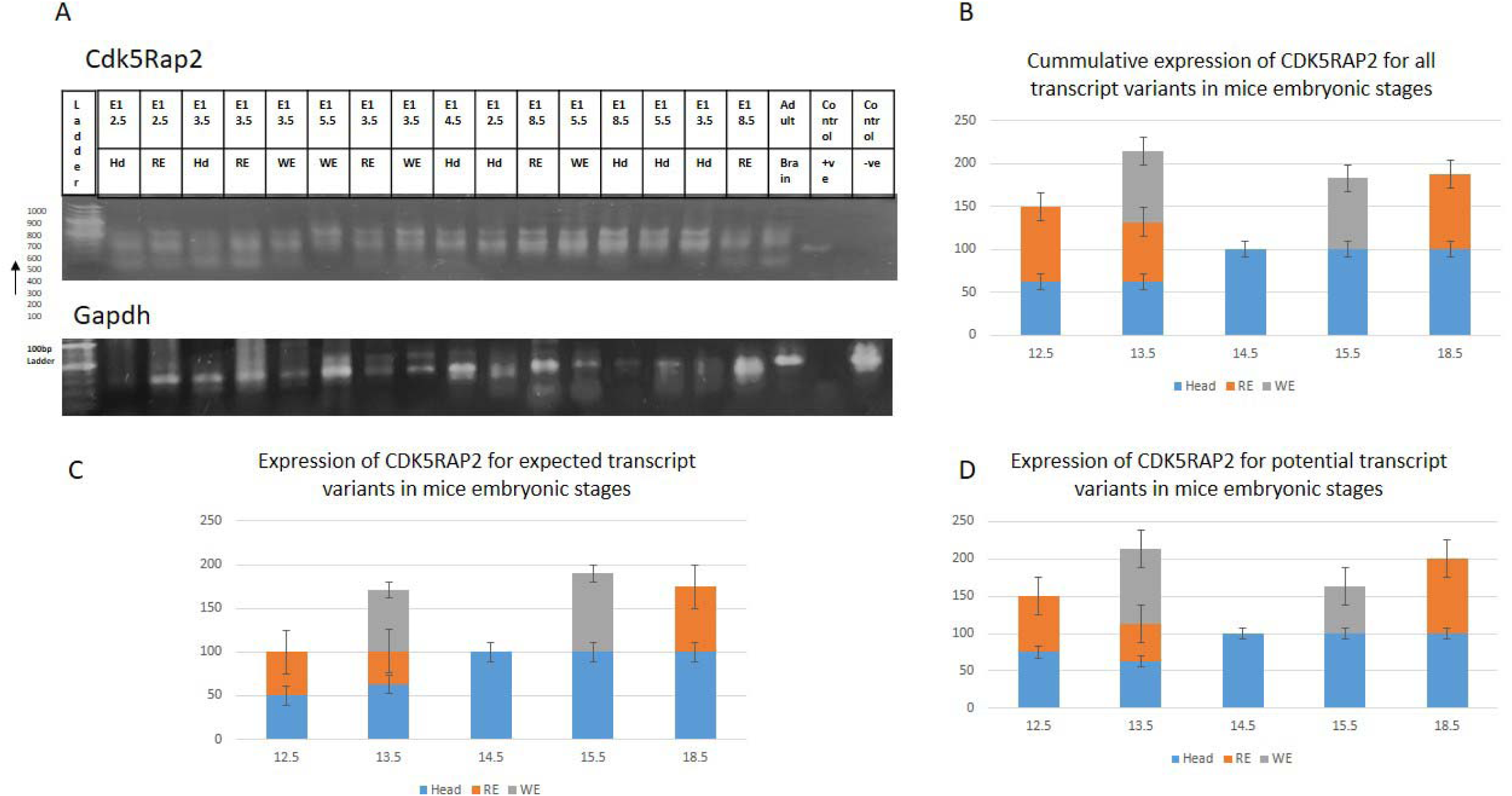
*Cdk5Rap2* an MCPH gene was subjected to transcriptional analysis by RT-PCR using gene specific primers. The *Gapdh* was used as a control. The results have shown expected band sizes and some potential new variants. The analysis was done by relative quantification of level of expression by band density and by given it a score from 4(100%) to 0(0%). The values for all different stages and embryonic regions were plotted in the graphs. A) Gel pictures, B) Cumulative expression at spatio-temporal dynamics C) Expected transcript variants scores D) New potential variants were plotted for each embryonic stage and region.

**Figure 3:**
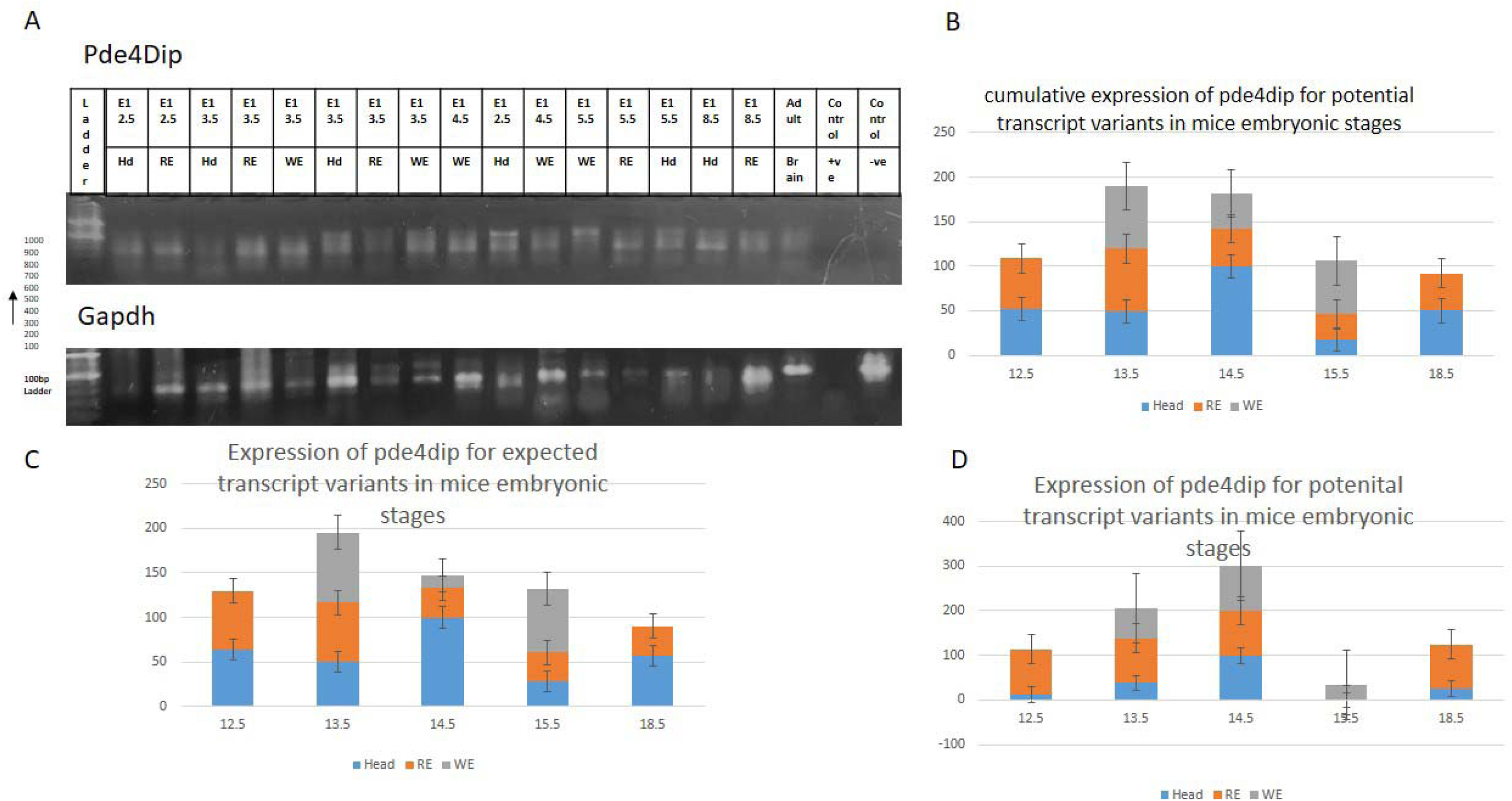
*Pde4dip/Myomegalin* was subjected to RT-PCR analysis with mRNA being by oligodT primers and dsDNA was made by gene specific primers. A) Gel pictures of *Pde4Dip* and *Gapdh* are shown at various embryonic days in mice in various embryonic regions (i.e., Head, rest of the embryo and whole embryos). B) The relative quantifications data for cumulative scores of all transcript variants is shown in various stages of neurogenesis in mice in various regions. C) Expected transcript variants data. D) Expression of potential new transcript variants.

Basal progenitors in humans and in the higher primates are the largest in number and contribute neurons to all six layers of the cerebral cortex and are thought to be the evolving factor between the highly convoluted gyrencephalic brain in higher mammals and the lissencephalic brain, as in the mice. Our data suggest that CDK5RAP2 and PDE4DIP play critical roles in these progenitors. Both *CDK5RAP2* and *PDE4DIP* are shown to be expressed in similar locations, such as centrosomes and involved in microtubule and spindle dynamics. Furthermore, the *PDE4DIP* if mutated cause anomalies of the cardiac system suggesting the divergent roles of these proteins in different organ development. We suggest that both of the proteins are essential for development where CDK5RAP2 specifically controls the brain development by prolonged expression throughout neurogenesis, whereas *PDE4DIP* is expressed in a small-time window that might be essential for the development of other organs.

### CEP63 and CCDC67

*CEP63* at SCKL6 on chromosome 3q22(Ma *et al*., 2014; Morris-Rosendahl and Kaindl, 2015; Sir *et al*., 2011a) is a gene involved in the broad development involving different organs and organ systems. The autosomal recessive mutation in *CEP63* gene causes an overall reduction, leading to dwarfism and severe microcephaly. The CEP63 protein is a core centrosomal protein involved in the centriole duplication, and recruitment of CEP152 at MCPH9 codes for a protein that also targets to the centrosome. Centriole duplication is an important component of the centrioler/centrosomal/spindle pole pathway, which is required for faithful mitosis and cell cycle progression. Its paralogue CCDC67/DEUP1(Zhao *et al*., 2013) is also studied to be involved in the centriole duplication and de novo centriole assembly and is therefore thought to complement the roles of CEP63 in development. *DEUP1* mutations have been reported to cause defects in the multiple-organs, in the affected individuals including retinopathy, cardiomyopathy, hypogonadism and others; however, a reduction in size of the brain has not been reported, and mental retardation is not common. Hence, although both proteins are important for the development, CEP63 seem to have brain-specific roles. The two proteins were subjected to PSI-BLAST, and MSA showed some smaller similar regions which were coiled coiled regions essential for centriole/centrosome targeting (see figure no. 4, A-C). We then started to investigate the spatio-temporal expression analysis of both the Seckel gene and the paralogue (see figure no. 5, A). We found that the cumulative expression of all the transcript variants for *Cep63* in the various embryonic samples at all stages of neurogenesis was more or less consistent from E12.5 to E18.5 with the highest expressions in E18.5 head (see figure no. 5, B). The novel transcript variants were expressed in the embryonic head at similar levels at E12.5, then down-regulated at E13.5 and then again up-regulated and remained at the same highest level as in E12.5, from E14.5 to E18.5 suggesting important functional relevance to the early and late progenitors (see figure no. 5, C-D). In contrast, *DEUP1* cumulative expression (see figure no. 6, A-D) was gradually up-regulated from E12.5 to E18.5, in a consistent way (see figure no. 6, B). The potential novel variants also showed the same trend, with gradual up-regulation from E12.5 to E18.5. The rest of the embryo, where we separated the embryonic head from the rest of the body, showed a gradual increase later from E15.5 to E18.5 (see figure no. 6, D). *Cep63* in the rest of the embryo samples also showed gradual up-regulation from E15.5 to E18.5, with the highest being at E18.5. The data suggests that the role of DEUP1 to be more general and is required in large copies at the end of neurogenesis and development, on the other hand, CEP63 seems to be highly essential in early to mid and late neurogenesis, with highest cumulative scores at E12.5, E15.5 and E18.5.

**Figure 4:**
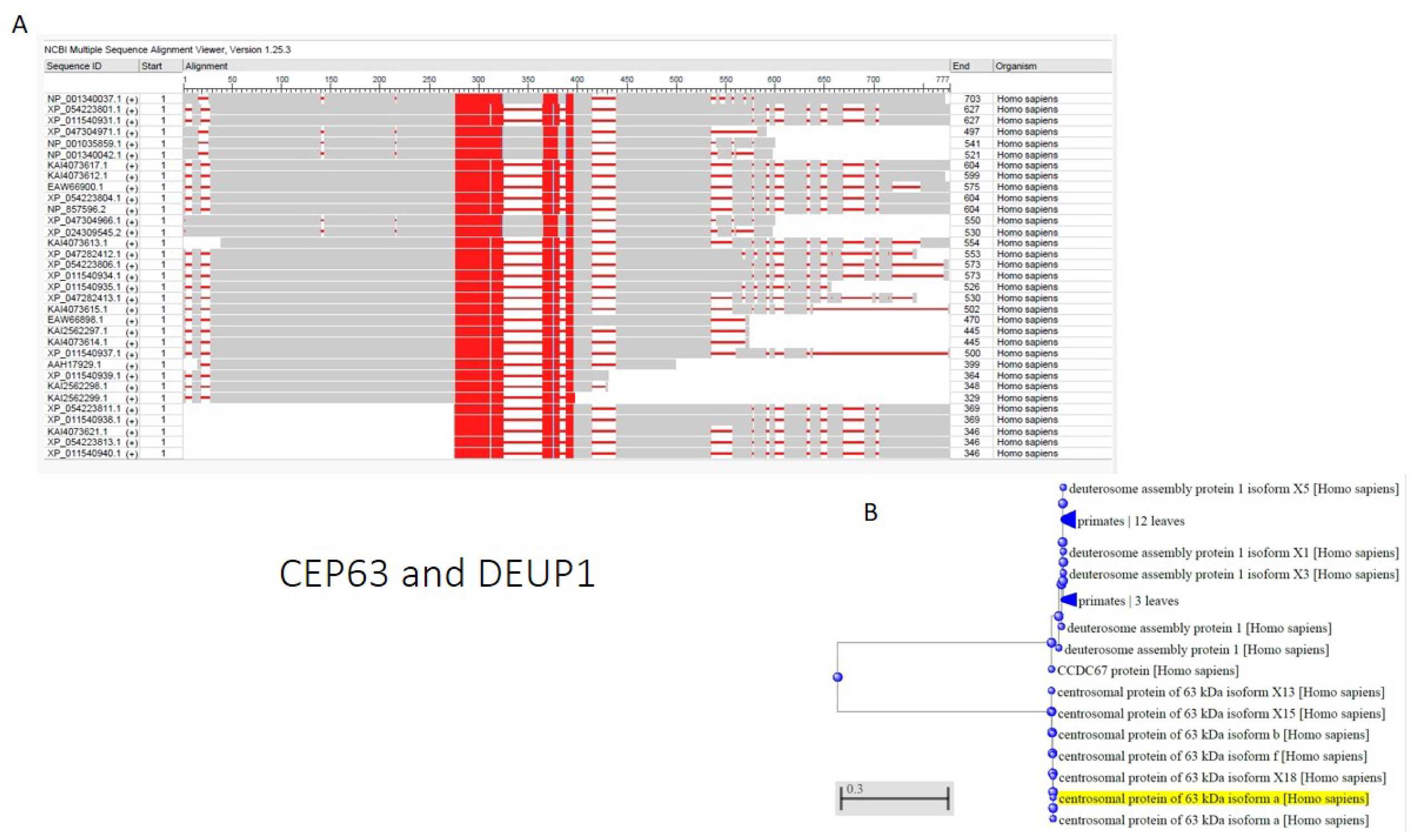
*CEP63* and *DEUP1/CCDC67* are two genes which are both undergoing active evolution and got split from a same ancestral gene. A) The Multiple Sequence Alignment of CEP63 is shown with various isoforms of DEUP1 and shows regions of high similarities scattered throughout but mostly in the center. Both proteins are targeted to the centrioles and the shared regions have coiled coiled domains important to the centrioler targeting. In addition to this, they also harbor a lot of varying regions where the two proteins are clearly showing divergence and new acquired functions B) Gene tree of both gene is shown where they have split in a single evolution event and undergoing active evolution.

**Figure 5:**
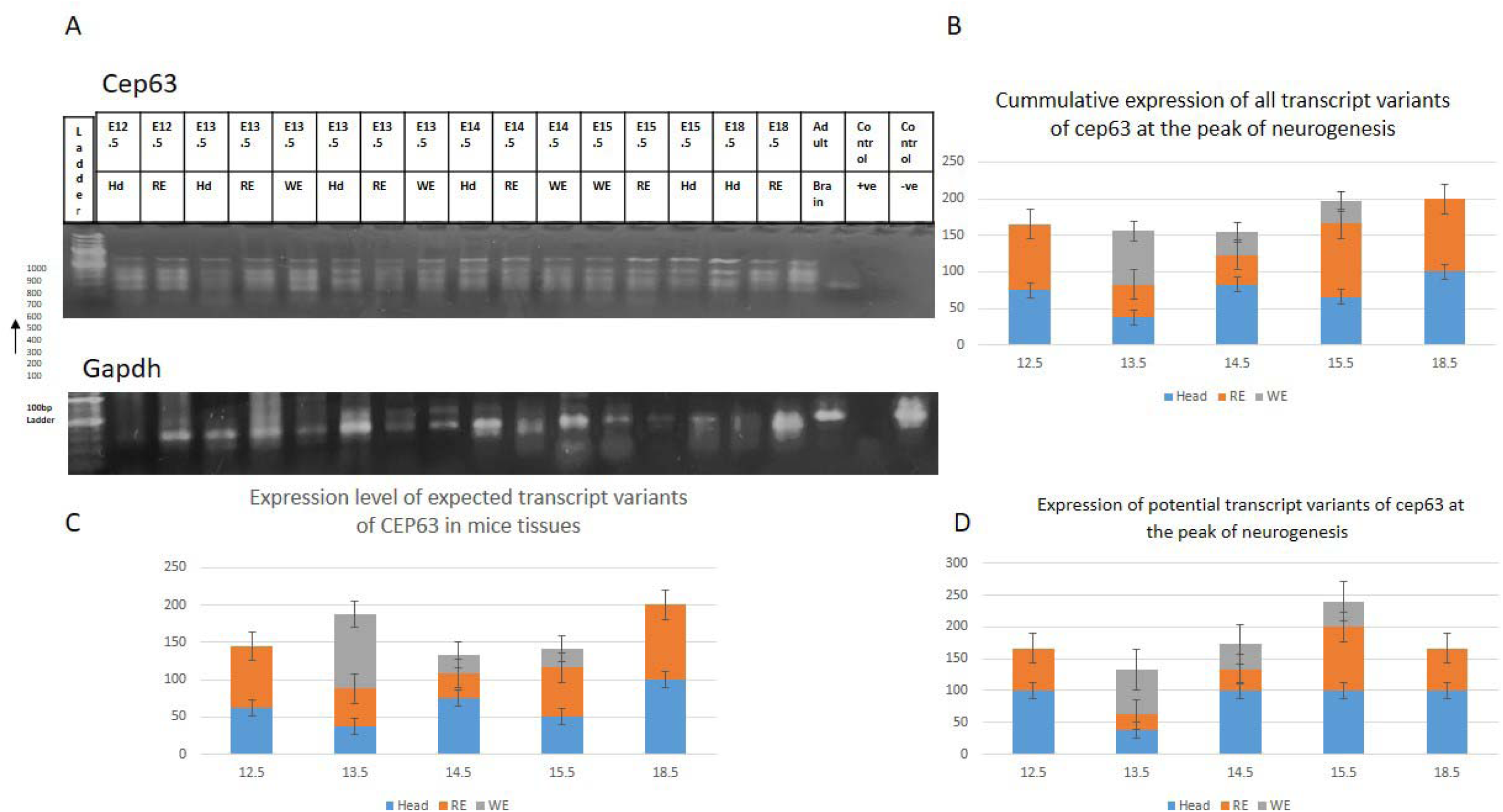
RT-PCR was done for *Cep63* throughout the peak of neurogenesis in mice i.e, E12.5 to E15.5 and E18.5. At each stage the mice embryos were divided in to the head regions, rest of the embryo and some whole embryos were also processed. Gene specific primers were utilized to do RT-PCR. A) Gel picture of RT-PCR in *Cep63* and Control *Gapdh* B) The cumulative expression of all the transcript variants showing highest expression at E12.5 but is showing quite consistent expression whereas head of E18.5 shows highest cumulative expression the time of OSVZ cell abundance. C) Expected transcript variants shows highest expression at E18.5 CD the potential new transcript variants shown expression at earlier and later stages.

**Figure 6:**
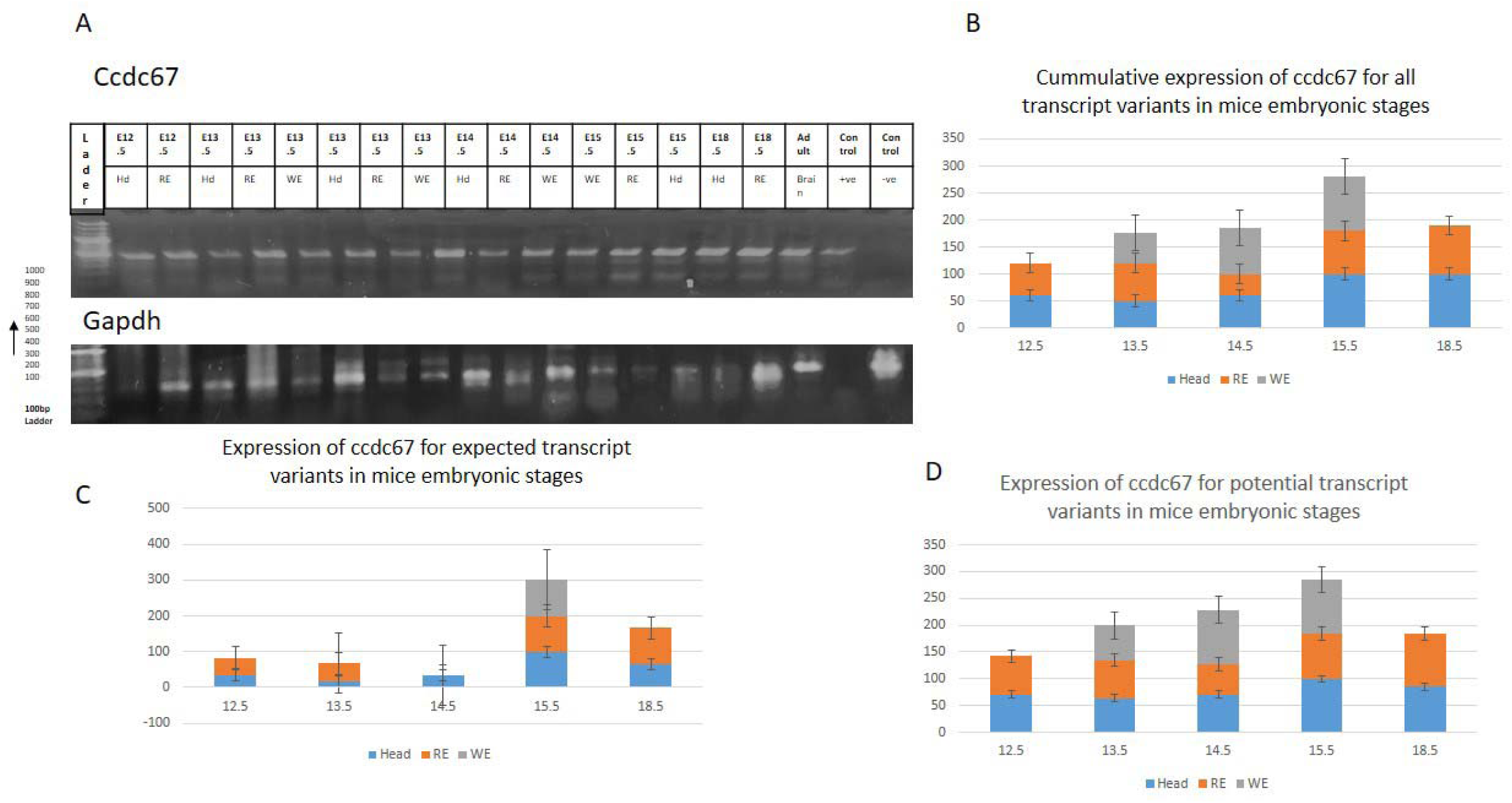
DEUP1 /CCDC67 a centriole targeted protein also a paralogue of Cep63 which also targets to centrioles and is a core component is subjected to transcriptional analysis by RT-PCR. A) Gels of *Ccdc67* and control *Gapdh* B) *Ccdc67* gradually increased from E12.5 to E18.5 reaching highest level gradually at later stages in cumulative scores C) The expected transcript variant also gradually increased D) The potential new variants showed same trend with quite consistent expression in the head region but collectively up-regulated from E12.5 to E18.5.

Both genes being highly up-regulated at mid to late gestation points shows their potential co-relation with the SVZ progenitors, which are abundant at these stages. There is also a small proportion of OSVZ progenitors in mice, which are detected in the late neurogenesis and are of high relevance in humans, and both genes seem to be also contributing to their neurogenesis. The rest of the embryo showed some upregulation and downregulation in *Cep63* but remained stable in Deup1 suggesting a more controlled and consistent expression of *Ccdc67*. These results suggest that at the time of development, Cep63 and Ccdc67 essentially control the brain development, but the other embryonic tissues require the maintenance of Ccdc67 throughout embryonic development. Hence, broad-spectrum disorders of the organs and organ systems are found in both *Cep63* as well as *Ccdc67* in mutated individuals.

### PHC1 and PHC3

*PHC1* and *PHC3* (Tonkin *et al*., 2002a) are the members of the Polyhomeotic-Like Proteins family, PLP, and are targeted to the PcG-PRC1, Polycomb group of repressor complexes, and are involved in the maintenance of pluripotency during development by repressing the transcription of genes involved in differentiation, especially *HOX* genes, in addition to other important roles in the nuclear dynamics (Awad *et al*., 2013a). This might be of significance in relation to MCPH, where a single change in PHC1 can lead to considerable brain size reduction, while leaving other organ systems intact. The *PHC3* is one of the paralogues of *PHC1* and was selected for this study because they split from the ancestral genes as two diverged genes in evolution as a single event and hence, could indicate separate acquired functions. The PSI-BLAST showed that the two paralogues are very similar and MSA showed C-terminus conservation, which are essential domains for chromatin remodeling by the PRC1 complex, suggestive of complementary roles (see figure no. 7 A-C). We set out to study their spatio-temporal dynamics by RT-PCR in mouse embryonic tissues at the times of neurogenesis from E12.5 to E18.5. RT-PCR results showed that the total expression of *Phc1* was highest at E12.5 and E13.5, with gradual downregulation at E14.5 and E15.5, whereas it again up-regulated at E18.5 (see figure no. 8, A-B). Expression in the embryonic head was the highest at E12.5, and later at E18.5. The expression of rest of embryonic tissues showed consistent expression from E12.5 to E18.5 (see figure no. 8, B-D). Hence indicating Phc1 to be essential for development in general and its controlled expression patterns in the brain indicated that it controls the early and late neurogenesis. Its paralogue, PHC3, which is thought to complement the function of Phc1 in the development, as being evolved from common ancestral gene. We then also investigated *Phc3* for mRNA expression for comparisons. Our results showed that *Phc3* was highly expressed at E12.5, slightly down regulated at E13.5 to E14.5, and enhanced at E15.5 to E18.5 being highest at these later stages (see figure no. 9 A& B). To dissect it further the embryonic heads and rest of embryos of mice showed consistent *Phc3* expression, which was highest at E12.5 in early neurogenesis, and later at E15.5 to E18.5 at late neurogenesis. The data indicate complementary roles for Phc1 and Phc3 in the brain, whereas early and late neurogenesis is controlled by these proteins (see figure no. 9, A-D). Interestingly, RT-PCR has showed a novel variant of Phc3 in embryonic heads of early embryos at E12.5 and E13.5 stage, in addition to the common variant which was expressed equally in all tissues, suggesting its relevance to the brain only (see figure no. 9, C). This suggests that in the brain, Phc1 function is complemented by Phc3 whereas in the rest of the embryos the Phc3 and Phc1 play essential diverged roles that might not be sensitive to single gene mutations.

**Figure 7:**
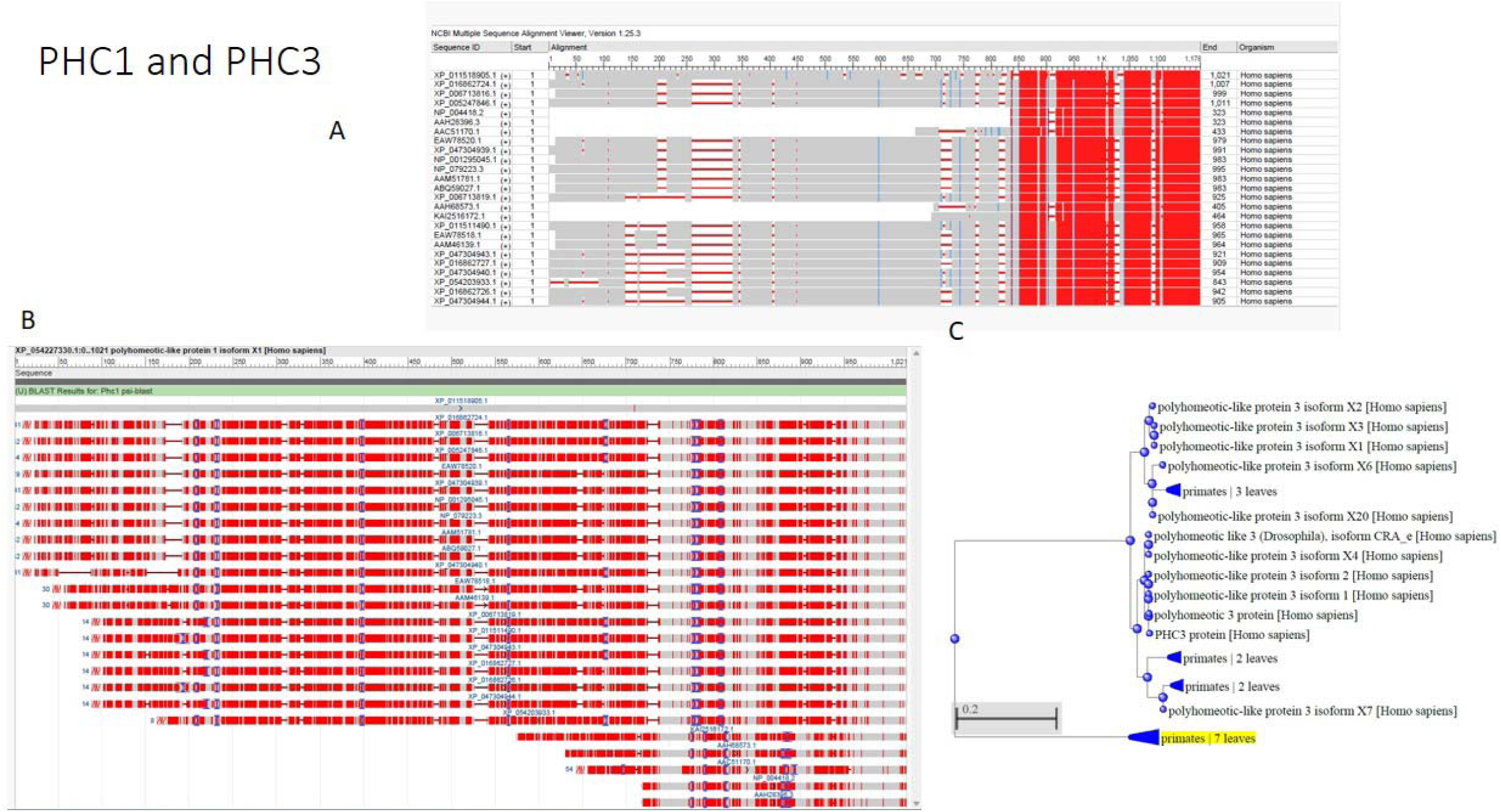
PSI-BLAST was used to identify, calculate similarity and ancestry of MCPH gene, *PHC1*’s paralogue, *PHC3*. Both genes are parts of PcG-PRC1 complex components and control pluripotency by chromatin remodeling. They seem to be targeted to the same pathway and were thought to be complementary to each other. A) PHC1 and PHC3 were aligned by Multiple Sequence Alignment and were shown to be highly similar in the C terminus regions and several other regions. The C terminus of both the proteins has FCS-Zinc finger domains and SAM domains essential for the chromatin remodeling B) The BLAST of Isoform 1 of PHC1 with all isoforms of PHC3 showing regions of similarity throughout in both proteins but very consistent C terminus C) The gene tree of *PHC1* and *PHC3* showing same ancestral gene and active evolution in PHC3.

**Figure 8:**
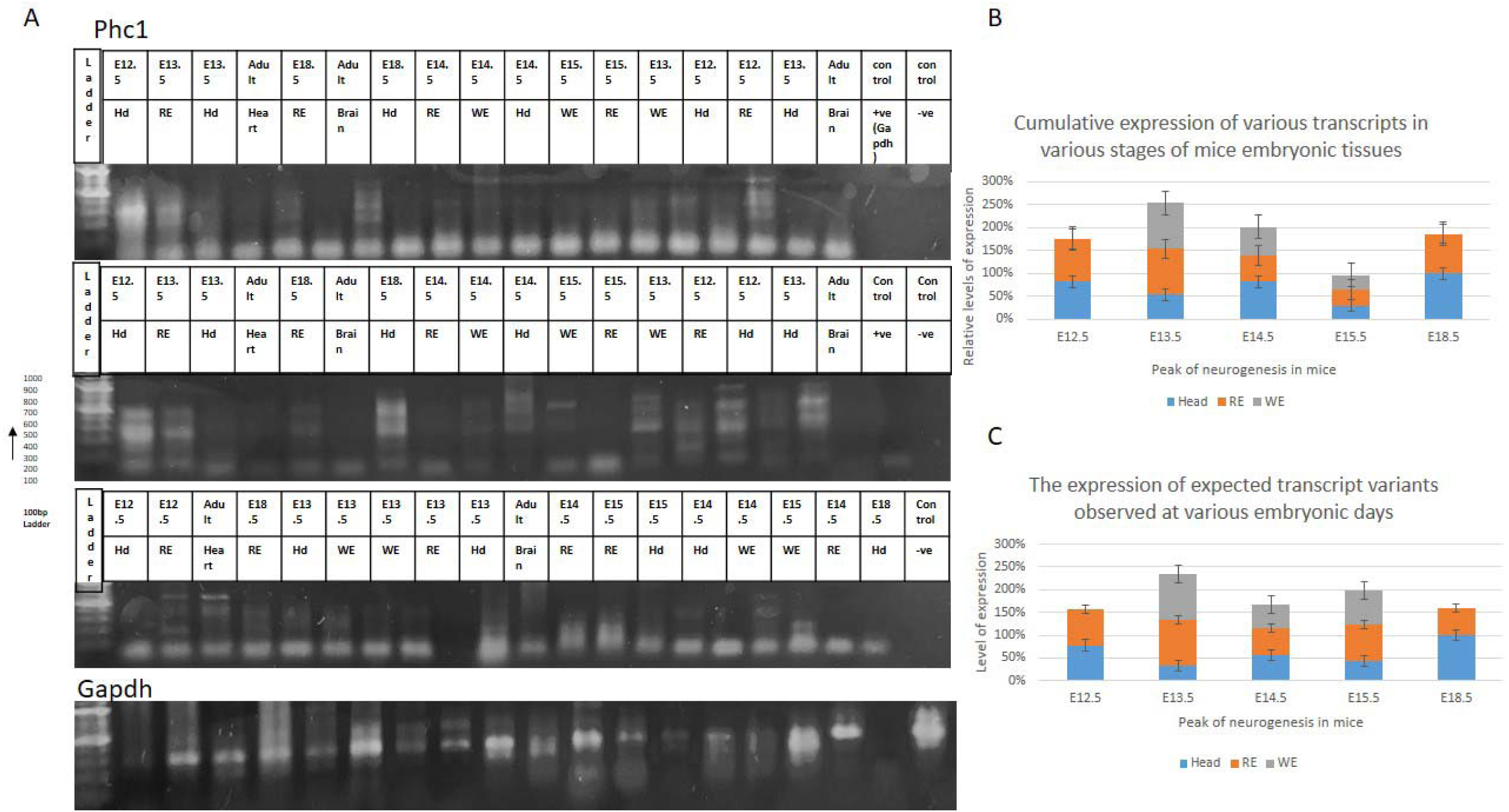
The RT-PCR data of Phc1 in various samples of mice embryos from peak of neurogenesis by gene specific primers. Three pairs of primers were used. A) Gel pictures of *Phc1* and Gapdh primers amplified products by ethidium bromide. B) Relative quantified band intensity for cumulative expression for all the transcript variants showing highest expression in the start, mid and end of neurogenesis but overall showing consistent expression which dips from E14.5 to E15.5 but again rises at E18.5 C) The potential new transcript shows similar trend but it relatively consistent at start and end.

**Figure 9:**
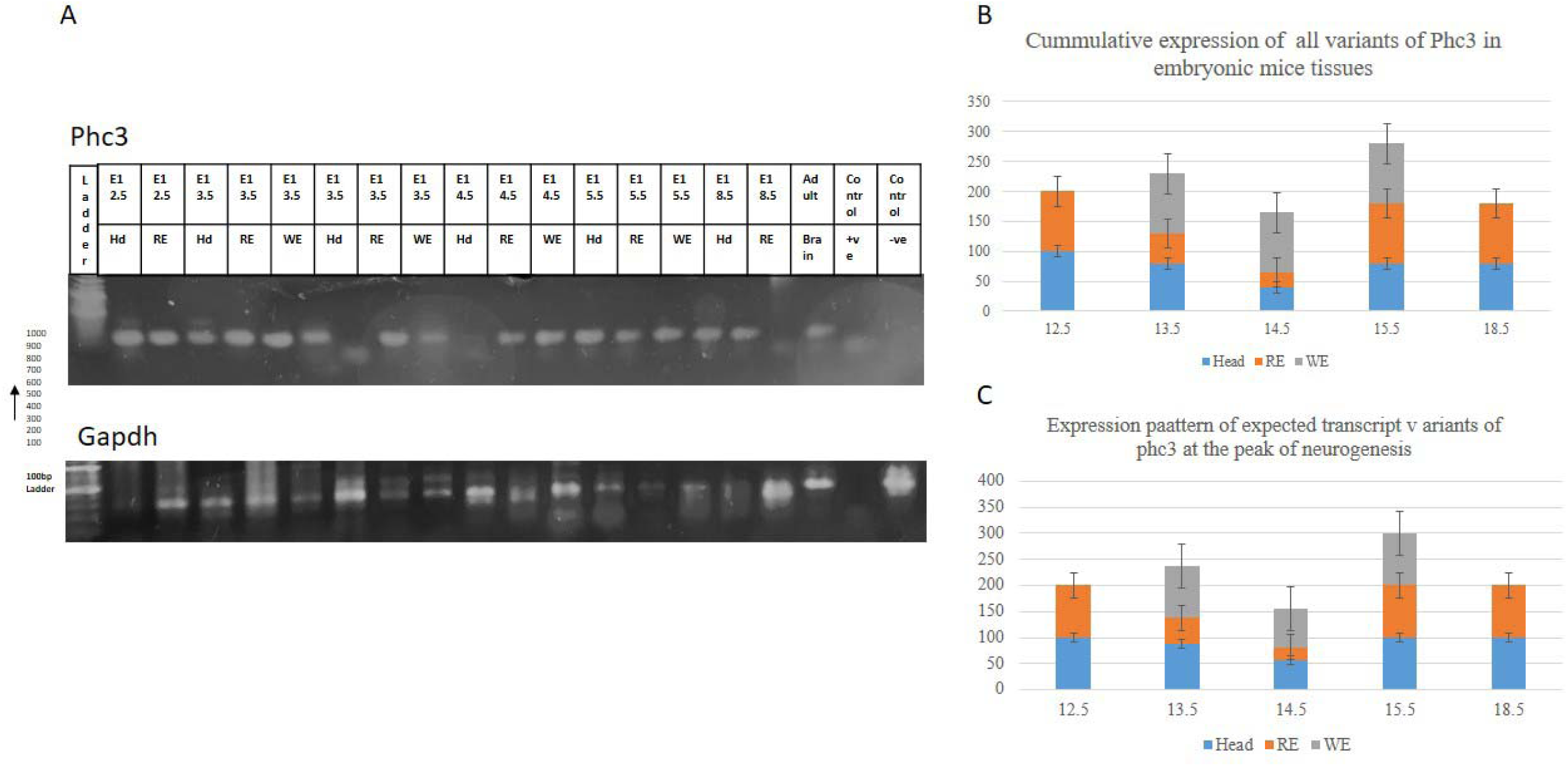
The RT-PCR data of PHC3 by gene specific primers. A) DNA Gel electrophoresis pictures of *Phc3* and control *Gapdh* showing the presence or absence and level of intensity of each sample bands. The peak of neurogenesis in mice was targeted by getting staged embryos from E12.5 to E15.5 and then E18.5 for late neurogenesis. The samples were prepared as head, Hd, rest of embryo, RE and Whole embryo, WE and are showing above each band B) The intensity level of each band was relatively quantified and given scores from 4(100%) to 0(0%) and were plotted for cumulative expression scores and found to be highest throughout neurogenesis with highest at earlier stage of E12.5 but otherwise consistent C) The potential new transcript variant was found to be present in E12.5 and E13.5 head and may be significant but all expected bands showed high trend in start and end of neurogenesis

During early and late neurogenesis, the radial glial cells and OSVZ cells are abundant and both have high importance in human brain development during neurogenesis. The radial glial cells being the initial progenitor cells that give rise to various progenitors and neuron lineages during neurogenesis, might be controlled by these two proteins in a way that PHC1 comes first and then PHC3 complements and control the final brain size. At MCP11 *PHC1*, if mutated leads to MCPH and indicate only brain reduction and other body organs are normal in size and might involve these spatio-temporal patterns.

#### Temporal dynamics of Genes and paralogues

Development is regulated by gene expression by transcription to RNA and then translation to proteins. Differential gene expression might be critical in order to regulate the normal development. Genetic neurodevelopmental disorders such as MCPH (microcephaly) and Seckel syndrome (microcephaly with dwarfism), seem to result from faulty genes, most of which reside in the centrosomes/spindle poles, but some also targeted to other compartments. Single gene mutations are causative agents of both, but with very different clinical profiles and overlapping diagnosis, i.e., primary microcephaly. We explored the idea of gene-paralogue pairs in both conditions by RT-PCR and did relative quantification and drew conclusions. Interestingly, however, when we explored the same data individually for each gene in the head of the embryo, Hd, and rest of embryo, RE (the embryonic body) tissues separately. We found the synchronized patterns in all three studied gene-paralogue pairs. In the mRNA of embryonic Hd tissues, one gene in each pair is having constant expression and the expression of other gene is going up and down with time showing increased and decreased expression. The synchronizing data is indicative of decisive proof of temporal dynamics during development in these tissues. Furthermore, the MCPH gene, *Cdk5Rap2* and *Pde4Dip* and Seckel gene, *Cep63* and *Ccdc67* pairs showed opposite patterns (see Figure 10, A-F) with the target gene being constant and the paralogue changing in the first case, whereas *Cep63* is continuously changing in expression with ups and down but paralogue is pretty much constant. Interestingly, the MCPH11 gene, *Phc1,* and its paralogue *Phc3* pair seem to show very similar profile as *Cep63-Ccdc67* with identical *Cep63* and *Phc1* patterns (Fig 10, B, C, E& F). This might be because the two genes are complementary and regulate brain development as a pair and seem to regulate this process by the continuous presence of one gene and changing behavior of the other gene in the pair which might be required for different neural progenitor types that arise as the neurogenesis proceeds. The genes were duplicated in evolution and may indicate the separate roles of each gene. We were further intrigued to study individual gene expression in the rest of the embryo, RE, in same time periods to see if there are any temporal changes in expression. It was again really fascinating, that we were seeing temporal changes with synchronization in patterns of different genes. This proves that not only temporally gene expression at mRNA level changing, but spatially it is also modulating. Interestingly, MCPH genes, seem to show synchronized patterns in the gene-paralogue pairs of *Cdk5Rap2-Pde4Dip* and *Phc1-Phc3* with *Phc1* and *Pde4Dip* being identical (see Figure 11, A-F). The *Phc1*’s paralogue, *Phc3* has same profile in Hd and RE (see Figure 10, F & 11, F) and seems to be required in same dosage throughout development. The Seckel pair, *Cep63* and *Ccdc67* seem to have the same patterns in the rest of the embryos and might indicate they can’t override each other during development and are required together. (see Figure 11 B, E).

**Figure no. 10:**
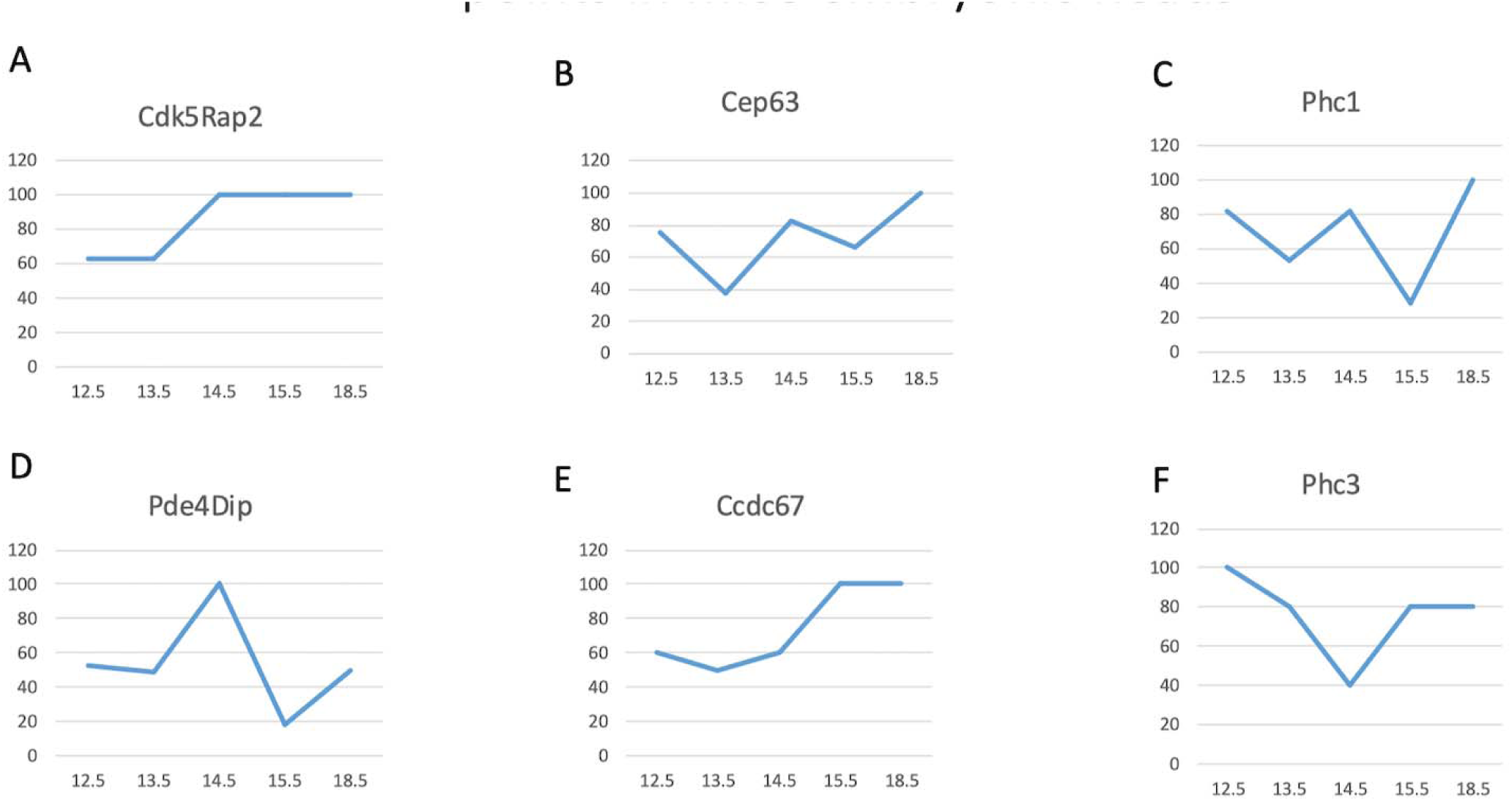
Temporal expression of gene-paralogue pairs in the heads of mice embryos at various embryonic days. The Cdk5Rap2 (A), Ccdc67(E) and Phc3(F) expression profiles were matching with their expression getting constant at later developmental stages in the head of embryos whereas Pde4Dip(D), Cep63(B) and Phc1(C) seem to have constantly changing and similar curves indicative of differential temporal gene expression.

**Figure no. 11:**
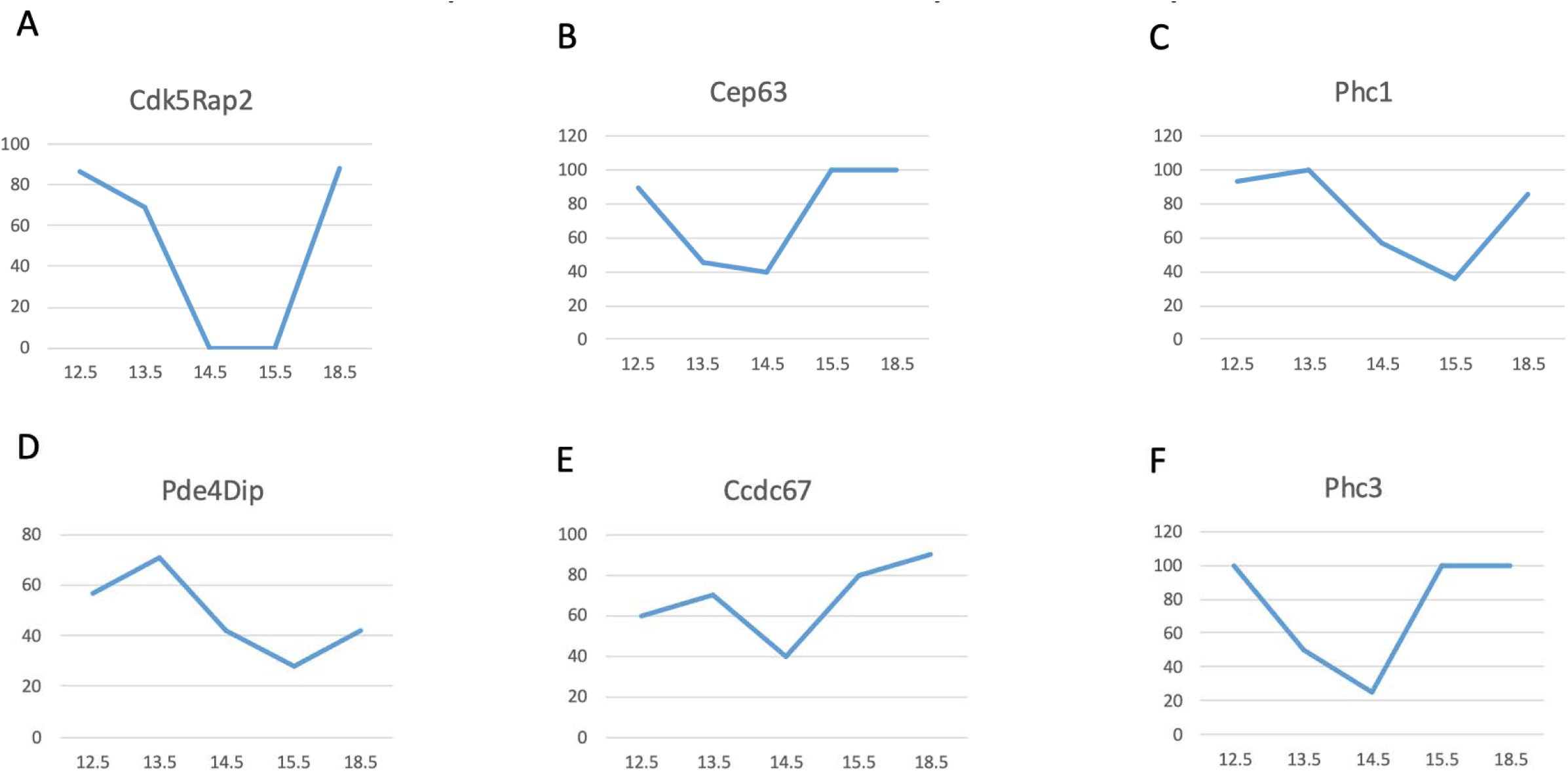
Temporal expression of genes and paralogues in the embryonic bodies (Rest of embryos, RE) of mice throughout neurogenesis. The Pde4Dip (D), Phc1(C) are identical in expression curves but total expression is different. The Cep63(B) Ccdc67(E) and Phc3(F) curves are matching and Cdk5Rap2(A) curve is not definitive due to two missing samples of RE at E14.5 and E15.5 and must be repeated.

#### The cumulative relative scores and total expression of each gene during neurogenesis

We then investigated whether these genes were sequentially upregulating or if they were required in a dose-dependent manner. We plotted the cumulative scores of each gene in stages of neurogenesis and found that in the Hd, the genes were at high level in the start of neurogenesis and at the later stages. In the RE, it is similar, but genes seem to have more constant gene profiles (see Figure 12, A & 13, A). The total expression of genes in both Hd and RE seems to suggest the same profile with *Phc3* being the most expressed among the six genes studied, and *Pde4Dip* was the least expressed (see Figure 12, B & 13, B). This can be viewed by keeping in mind the idea of a single master pathway where, sequentially, the genes are upregulated, but another possibility also exists that these genes might have differing roles in development and different doses of each are required or both. Furthermore, in the RE, the total expression profile of all three genes, *Cdk5Rap2*, *Cep63* and *Phc1* are more or less balanced with one paralogue, *Phc3* being the highest and at a similar level to its paralogue, *Phc1* (see Figure 13, B). This re-iterates our earlier suggestion that the microcephaly gene, *Phc1* and its paralogue, *Phc3*, are required for development together and are complementing each other and are targeted to the same pathway. In Hd tissues, the changing doses of genes are required in order to control the division of various progenitors which are brain specific only and hence control brain development through temporal dynamics (see Figure 12, A-B). The other two gene-paralogue pairs, have seem to be divergent and control different functions during development especially the Seckel gene-paralogue pair.

**Figure no. 12:**
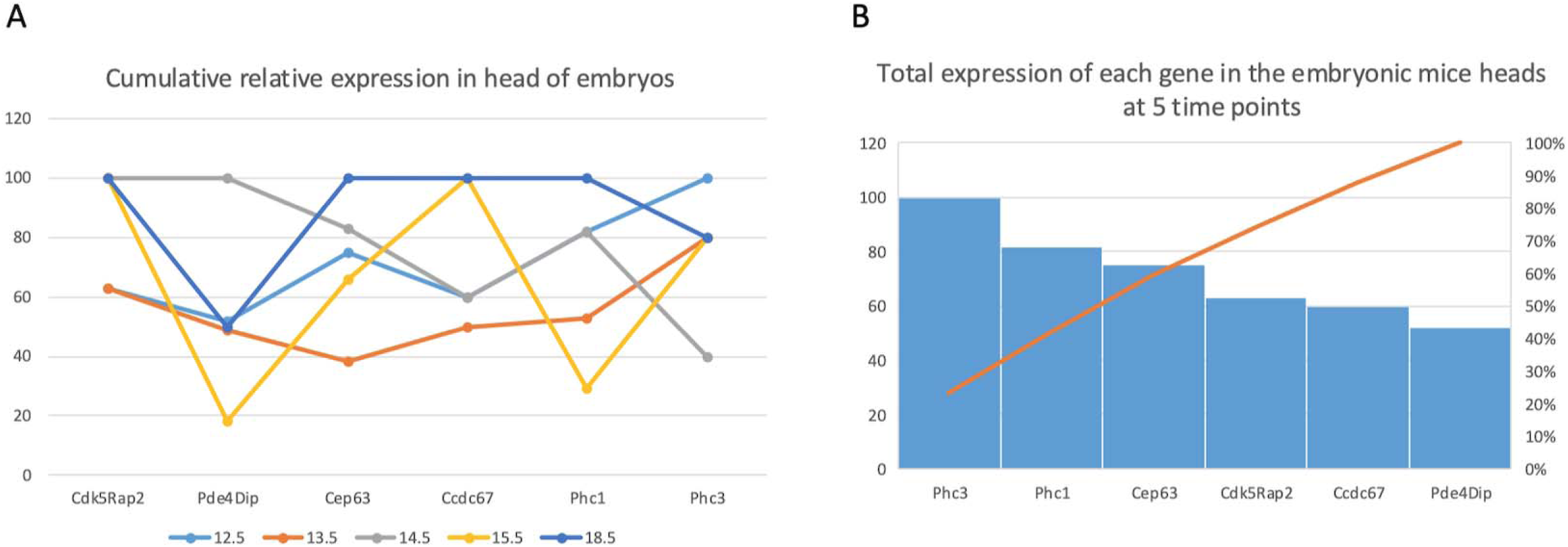
Relative quantification of all six genes at various time points throughout neurogenesis (Temporal expression). A) Cumulative relative expression of genes in embryonic heads. B) Total expression of each gene in the embryonic heads in mice at E12.5, E13.5, E14.5, E15.5 and E18.5 and shows a linear curve.

**Figure no. 13:**
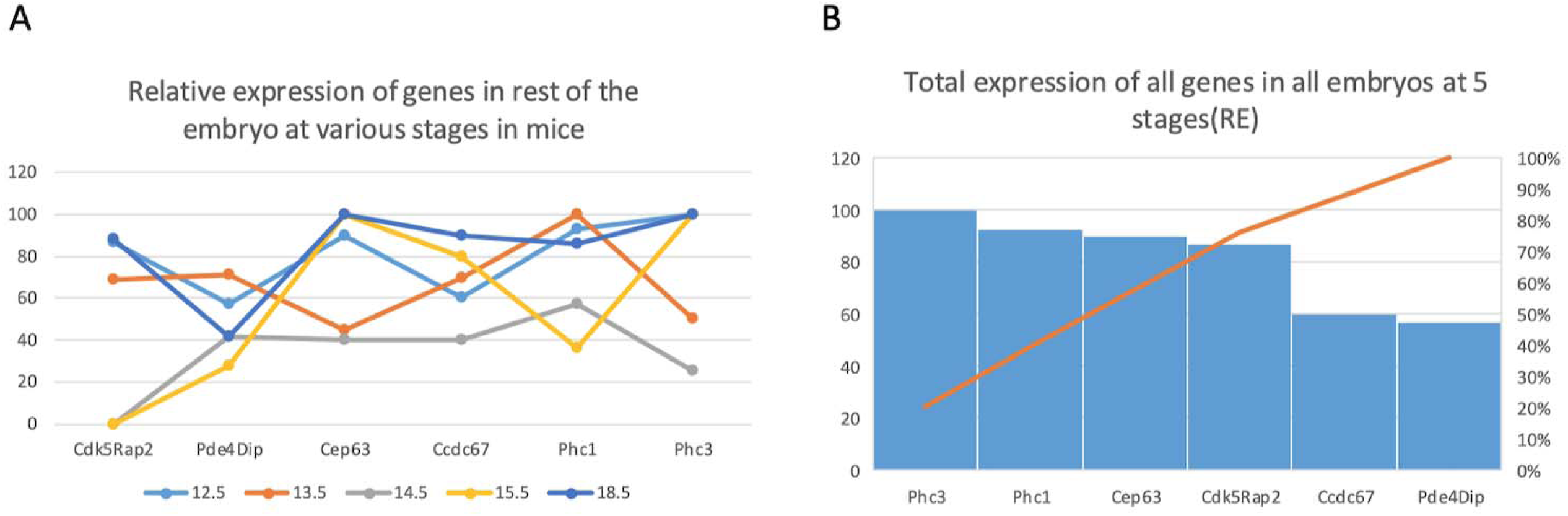
Relative quantification of all six genes at various time points throughout neurogenesis (Temporal expression). A) Cumulative relative expression of genes in embryonic bodies, RE. B) Total expression of each gene in the embryonic bodies, RE, in mice at E12.5, E13.5, E14.5, E15.5 and E18.5 and shows a linear curve.

## Discussion

MCPH and Seckel are disorders involving a very tragic microcephaly condition which is further added upon by dwarfism in the syndromic later form. Scientists are also fascinated by this condition because the microcephaly is similar to the head sizes in our closest relatives, the higher primates. Not only the size of the head, which is most notably small in the cerebral cortex region that is exceptionally developed in *Homo Sapiens,* but it also limits the children to learn language or perform very complex tasks and moderately to severe mental retardation, which might just reflect their inability to mimic human-specific brain functions but are quite decent. It is a disorder that opens up new avenues for studying evolution and human-acquired traits. Most scientific work in the past was focused on the discovery of the pathogenic genes, which was challenging and time-consuming, and this was accelerated later by the advent of high throughput sequencing and computational analysis (Asif *et al*., 2023b; Kaindl *et al*., 2010b; Thornton and Woods, 2009b; Woods, 2004; Zaqout and Kaindl, 2022b) In addition to this, some very important functional analysis was done on these genes and found to be highly informative (Awad *et al*., 2013b; Bond *et al*., 2005; Dell’amico *et al*., 2023; Jayaraman *et al*., 2016b, 2016b; Kouprina *et al*., 2005; Marthiens and Basto, 2014; Nicholas *et al*., 2010b; Rickmyre *et al*., 2007; Sir *et al*., 2011b; Yang *et al*., 2012), but gave little insight in to the mode of pathogenesis of the disorder. All of these genes are part of essential core pathways that control cells and neural progenitors as the type of cell also have it in all cells and seem to be ubiquitous more or less (Florio and Huttner, 2014; Huttner *et al*., 2006). Furthermore, their upregulation or downregulation has also been reported in the many cancers and tumor cells, indicating their regulated roles. Patient-derived cell lines show reduced or absent expression, but this is also not a true reflection when we think of it as a developmental disorder. Animal models have shown some promise, but with varying results (Brunk *et al*., 2007; Kronman *et al*., 2024; Pilz *et al*., 2018; Pulvers and Huttner, 2009; Rickmyre *et al*., 2007; Tonkin *et al*., 2002b; Wainman *et al*., 2009). We set out to the basic developmental principal, where genes are differentially regulated with respect to stage and time and control different specific roles that they play in the development (Duan *et al*., 2022b; Georgala *et al*., 2011; Mannino *et al*., 2023; Roberson *et al*., 2021; Vaid and Huttner, 2020b). We set out to study three pairs of genes and their paralogues which are targeted to different pathways and are involved in MCPH and/or Seckel syndrome. We used RT-PCR to check if there was any variation in their expression levels or presence or absence of transcripts at various stages throughout the peak of neurogenesis. We numerically calculated the levels of presence or absence and sometimes used many primers targeted to different exons and added up all scores for each band and calculated the relative intensity scores. Interestingly these have actually provided us with positive insight in to the disorders; this is the first report to actually do so. This will further open new chapters to exciting research on DNA/RNA and protein levels in the future and combining high-tech machines could provide important insights in to this field of spatio-temporal dynamics (Duan *et al*., 2022b; Kim and An, 2020; Roberson *et al*., 2021; West *et al*., 2022) during development.

Our data suggests that neural progenitor cell divisions into neurons through symmetric and asymmetric divisions are controlled by various cellular pathways, and MCPH genes indicate a series of upregulation at different embryonic days suggesting their overall hierarchy in the control of neurogenesis. In addition, their respective paralogues seem to complement their roles in neural development which might indicate their divergence in the development of other organs apart from the brain. *Cdk5Rap2-Pde4dip* showed up-regulation at mid-late neurogenesis and seem to have some redundancy whereas *Cep63-Ccdc67* duo was very different and expression of *Cep63* was consistent throughout neurogenesis, but *Ccdc67* gradually increased during development and was highest at the late neurogenesis and development suggestive of diverged and essential roles of the two genes which are both essential in development and lastly, *Phc1-Phc3* showed up-regulation at early and late neurogenesis and might be complimentary for brain development. This suggests the roles of different pathways are important for cell divisions at different times and stages, and might be possibly part of a higher single master pathway that incorporates all the mini-pathways. All the genes studied showed spatiotemporal(Pang *et al*., 2024) variation in expression and when plotted on their own in terma of temporal expression change at the studied stages in two compartments, embryonic heads, Hd, and embryonic body, RE. The genes seem to distribute in two common pairs of similar patterns of temporal expression changes consisting of actively varying in *Pde4Dip*, *Cep63* and Phc1 and of more increased expression at earlier stages and constant in later stages in *Cdk5Rap2*, *Ccdc67* and *Phc3* which seem to suggest that these gene-paralogue duos are actually similar and result in neurodevelopmental disorders if any is mutated. The *Cep63* is Seckel syndrome causing gene and is more severe condition with overall reduction in body size and other ailments so we were interested to look at the individual gene profiles in the RE samples from embryos and again data was really interesting regarding grouping of genes in two categories. *Pde4Dip* and *Phc1* had same profile and was actively changing and was also upregulating later at these stages whereas, *Cep63* and *Phc3* was same. *Cdk5Rap2* cannot be fully recapitulated due to two missing sample data for RE at E13.5 and E14.5 which may be repeated in future. *Ccdc67* was similar to *Cep63* and seem to suggest both proteins to be essential for body development and reason that Seckel syndrome results for missing Cep63. These dynamics need to be studied for other MCPH and Seckel genes and their paralogues, and the mRNA data must be validated by protein expression analysis, and the work may also be replicated in human-based models/ fetal tissuesor single cells from mice (Brooks et al., 2024). The mRNA has been shown to transcribe in bursts and this phenomenon may also be studied(Pomp *et al*., 2024) and our results may also be due to this phenomenon of ups and downs with time but may be regulated differently due to unique nature of developing brain cells. The special developing brain cells need these differential cues to changing population of stem and progenitor cells having different fate and differentiation potential and these genes might be important for brain and overall development and might regulate both by complementing each other or by being having different controls.

## Future prospects

The study is only preliminary and does not fully recapitulate all the MCPH and Seckel genes, which exceed to more than 45. It is also required that these data should be verified by total gene expression variation in each tissue and stage by qRT-PCR. Furthermore, single-cell expression profiling must be performed to obtain definitive answers to spatio-temporal changes. High-throughput mRNA and protein analysis of these genes and paralogues must be done on single cell samples in order to determine the neural stem and progenitor cells for differential expression and then deduce and relate it to primary microcephaly (Birk *et al*., 2024; Winkler *et al*., 2024). Human brain organoids may be able to answer these questions at the beginning.

## Acknowledgements

We want to thank all members of our lab for their input in the form of work support and team work.

## Data and resource availability

The study used gene-specific primers and is included as supplementary data. Furthermore, a preprint of the paper with data is on open preprint server, (Mohsin *et al*., 2025). Other supporting data if required will be made available.

## Diversity and inclusion statement

We are do not discriminate anyone on the basis of race, nationality, religion or gender.

## Authors contributions

M.M., S.R., did the investigation and data curation, M.M, also helped with formal analysis, M.I., did supervision, J.J.C., helped with writing and agreed to editing of the manuscript post acceptance and M.K. visualized the project, secured funding, did project administration, planned methodology, did formal analysis, did validation and wrote the manuscript.

## Supplementary data

The supporting data in form of calculation and primers sequence are provided with the article as S1 Table and S2 Table.

