## Supplementary material for "The study of differential expressions of MCPH and Seckel syndrome genes and their paralogues": S1 Table, S2 Table

**Supplementary Data tables paralogue paper**

**S1 Table - Relative quantitation**

| ***CDK5RAP2*** | | | **Expected 754 bp, 973bp** | | | **Observed 973 754 400bp** | | | |
| --- | --- | --- | --- | --- | --- | --- | --- | --- | --- |
| **Cumulative** | | | | | | | | | |
| **Embryonic Day** | **Head** | **Total intensity** | **% relative expression** | **RE** | **Total intensity** | **% relative expression** | **WE** | **Total intensity** | **% relative expression** |
| 12.5 | 0,1,2 | 3 | 38 | 0,2,3 | 5 | 87 | Sample absent |  |  |
| 12.5 | 0,3,4 | 7 | 87 | Sample absent |  |  | Sample absent |  |  |
| 13.5 | 0,1,1 | 2 | 25 | 0,1,4 | 5 | 87 | 0,2,4 | 6 | 66 |
| 13.5 | 0,4,4 | 8 | 100 | 0,2,2 | 4 | 50 | 1,4,4 | 9 | 100 |
| 14.5 | 0,4,4 | 8 | 100 | Sample absent |  |  | Sample absent |  |  |
| 15.5 | Sample absent |  |  | Sample absent |  |  | 1,4,2 | 7 | 77 |
| 15.5 | 0,4,4 | 8 | 100 |  |  |  | 0,4,4 | 8 | 88 |
| 18.5 | 0,4,4 | 8 | 100 | 0,2,4 | 6 | 75 |  |  |  |
| 18.5 | Sample absent |  |  | 0,4,4 | 8 | 100 | Sample absent |  |  |
| **Potential new variant** | | | | | | | | | |
| **Embryonic Day** | **Head** | **Total intensity** | **% relative expression** | **RE** | **Total intensity** | **% relative expression** | **WE** | **Total intensity** | **% relative expression** |
| 12.5 | 2 | 2 | 50 | 3 | 3 | 75 | Sample absent |  |  |
| 12.5 | 4 | 4 | 100 | Sample absent |  |  | Sample absent |  |  |
| 13.5 | 1 | 1 | 25 | 2 | 2 | 50 | 4 | 4 | 100 |
| 13.5 | 4 | 4 | 100 | 2 | 2 | 50 | 4 | 4 | 100 |
| 14.5 | 4 | 4 | 100 | Sample absent |  |  | Sample absent |  |  |
| 15.5 | Sample absent |  |  | Sample absent |  |  | 2 | 2 | 50 |
| 15.5 | 4 | 4 | 100 |  |  |  | 4 | 4 | 100 |
| 18.5 | 4 | 4 | 100 | 4 | 4 | 100 | Sample absent |  |  |
| 18.5 | Sample absent |  |  | 4 | 4 | 100 | Sample absent |  |  |
| **Expected Variants** | | | | | | | | | |
| **Embryonic Day** | **Head** | **Total intensity** | **% relative expression** | **RE** | **Total intensity** | **% relative expression** | **WE** | **Total intensity** | **% relative expression** |
| 12.5 | 0,1 | 1 | 25 | 0,2 | 2 | 50 | Sample absent |  |  |
| 12.5 | 0,3 | 3 | 75 | Sample absent |  |  | Sample absent |  |  |
| 13.5 | 0,1 | 1 | 25 | 0,1 | 1 | 25 | 0,2 | 2 | 40 |
| 13.5 | 0,4 | 4 | 100 | 0,2 | 2 | 50 | 1,4 | 5 | 100 |
| 14.5 | 0,4 | 4 | 100 | Sample absent |  |  | Sample absent |  |  |
| 15.5 | Sample absent |  |  | Sample absent |  |  | 1,4 | 5 | 100 |
| 15.5 | 0,4 | 4 | 100 | Sample absent |  |  | 0,4 | 4 | 80 |
| 18.5 | 0,4 | 4 | 100 | 0,2 | 2 | 50 | Sample absent |  |  |
| 18.5 | Sample absent |  |  | 0,4 | 4 | 100 | Sample absent |  |  |

| ***PDE4DIP*** | | | **Expected 252, 405bp** | | | | **Observed 520,405,252bp** | | |
| --- | --- | --- | --- | --- | --- | --- | --- | --- | --- |
| **Cumulative** | | | | | | | | | |
| **Embryonic Day** | **Head** | **Total intensity** | **% relative expression** | **RE** | **Total intensity** | **% relative expression** | **WE** | **Total intensity** | **% relative expression** |
| 12.5 | 1,1,3 | 5 | 50 | 0,1,3 | 4 | 57 | Sample absent |  |  |
| 12.5 | 0,2,4 | 6 | 54 | Sample absent |  |  | Sample absent |  |  |
| 13.5 | 0,0,1 | 1 | 16 | 1,3,3 | 7 | 100 | 1,1,3 | 4 | 40 |
| 13.5 | 3,3,3 | 9 | 81 | 1,1,1 | 3 | 42 | 3,3,4 | 10 | 100 |
| 14.5 | 4,3,4 | 11 | 100 | 1,1,1 | 3 | 42 | 3,1,0 | 4 | 40 |
| 15.5 | 0,0,2 | 2 | 18 | 0,0,2 | 2 | 28 | 1,1,4 | 6 | 60 |
| 18.5 | 1,2,2 | 5 | 50 | 1,1,1 | 3 | 42 | Sample absent |  |  |
| **Potential new variants** | | | | | | | | | |
| **Embryonic Day** | **Head** | **Total intensity** | **% relative expression** | **RE** | **Total intensity** | **% relative expression** | **WE** | **Total intensity** | **% relative expression** |
| 12.5 | 1, | 1 | 25 | 1 | 1 | 100 | Sample absent |  |  |
| 12.5 | 0 | 0 | 0 | Sample absent |  |  | Sample absent |  |  |
| 13.5 | 0 | 0 | 0 | 1 | 1 | 100 | 1 | 1 | 33 |
| 13.5 | 3 | 3 | 75 | 1 | 1 | 100 | 3 | 3 | 100 |
| 14.5 | 4 | 4 | 100 | 1 | 1 | 100 | 3 | 3 | 100 |
| 15.5 | 0 | 0 | 0 | 0 | 0 |  | 1 | 1 | 33 |
| 18.5 | 1 | 1 | 25 | 1 | 1 | 100 | Sample absent |  |  |
| **Expected transcript variants** | | | | | | | | | |
| **Embryonic Day** | **Head** | **Total intensity** | **% relative expression** | **RE** | **Total intensity** | **% relative expression** | **WE** | **Total intensity** | **% relative expression** |
| 12.5 | 1,2 | 3 | 42 | 1,3 | 4 | 66 | Sample absent |  |  |
| 12.5 | 2,4 | 6 | 85 | Sample absent |  |  | Sample absent |  |  |
| 13.5 | 0,1 | 1 | 14 | 3,3 | 6 | 100 | 1,3 | 4 | 57 |
| 13.5 | 3,3 | 6 | 85 | 1,1 | 2 | 33 | 3,4 | 7 | 100 |
| 14.5 | 3,4 | 7 | 100 | 1,1 | 2 | 33 | 1,0 | 1 | 14 |
| 15.5 | 0,2 | 2 | 28 | 0,2 | 2 | 33 | 1,4 | 5 | 71 |
| 18.5 | 2,2 | 4 | 57 | 1,1 | 2 | 33 | Sample absent |  |  |

| ***CEP63*** | | | **Expected 480bp, 302bp** | | | **Observed 700, 480, 302, 180bp** | | | |
| --- | --- | --- | --- | --- | --- | --- | --- | --- | --- |
| **Cumulative** | | | | | | | | | |
| **Embryonic Day** | **Head** | **Total intensity** | **% relative expression** | **RE** | **Total intensity** | **% relative expression** | **WE** | **Total intensity** | **% relative expression** |
| 12.5 | 1,2,3,3 | 9 | 75 | 1,2,3,3 | 9 | 90 | Sample absent |  |  |
| 13.5 | 1,1,1,1 | 4 | 33 | 1,2,2,1 | 6 | 60 | 2,4,4,3 | 13 | 100 |
| 13.5 | 1,2,2,0 | 5 | 42 | 1,1,1,0 | 3 | 30 | 1,2,2,1 | 6 | 46 |
| 14.5 | 2,3,3,2 | 10 | 83 | 1,1,1,1 | 4 | 40 | 1,1,1,1 | 4 | 31 |
| 15.5 | 4,2,2,0 | 8 | 66 | 4,2,2,2 | 10 | 100 | 1,1,1,1 | 4 | 30 |
| 18.5 | 4,4,4,0 | 12 | 100 | 3,3,3,1 | 10 | 100 | 2,4,4,1 | 11 | 84 |
| **Potential new variant** | | | | | | | | | |
| **Embryonic Day** | **Head** | **Total intensity** | **% relative expression** | **RE** | **Total intensity** | **% relative expression** | **WE** | **Total intensity** | **% relative expression** |
| 12.5 | 1,3 | 4 | 100 | 1,3 | 4 | 66 | Sample absent |  |  |
| 13.5 | 1,1 | 2 | 50 | 1,1 | 2 | 33 | 2,3 | 5 | 100 |
| 13.5 | 1,0 | 1 | 25 | 1,0 | 1 | 16 | 1,1 | 2 | 40 |
| 14.5 | 2,2 | 4 | 100 | 1,1 | 2 | 33 | 1,1 | 2 | 40 |
| 15.5 | 4,0 | 4 | 100 | 4,2 | 6 | 100 | 1,1 | 2 | 40 |
| 18.5 | 4,0 | 4 | 100 | 3,1 | 4 | 66 | 2,1 | 3 | 60 |
| **Expected transcript variants** | | | | | | | | | |
| **Embryonic Day** | **Head** | **Total intensity** | **% relative expression** | **RE** | **Total intensity** | **% relative expression** | **WE** | **Total intensity** | **% relative expression** |
| 12.5 | 2,3, | 5 | 62 | 2,3 | 5 | 83 | Sample absent |  |  |
| 13.5 | 1,1 | 2 | 25 | 2,2 | 4 | 66 | 4,4 | 8 | 100 |
| 13.5 | 2,2, | 4 | 50 | 1,1 | 2 | 33 | 2,2 | 4 | 50 |
| 14.5 | 3,3 | 6 | 75 | 1,1 | 2 | 33 | 1,1 | 2 | 25 |
| 15.5 | 2,2 | 4 | 50 | 2,2 | 4 | 66 | 1,1 | 2 | 25 |
| 18.5 | 4,4 | 8 | 100 | 3,3 | 6 | 100 | 4,4 | 8 | 100 |

| ***Ccdc67*** | | | **Expected**  **423bp** | | | | **observed**  **500, 423, 190bp** | | |
| --- | --- | --- | --- | --- | --- | --- | --- | --- | --- |
| **Cumulative** | | | | | | | | | |
| **Embryonic Day** | **Head** | **Total intensity** | **% relative expression** | **RE** | **Total intensity** | **% relative expression** | **WE** | **Total intensity** | **% relative expression** |
| 12.5 | 4,1,1 | 6 | 60 | 4,1,1 | 6 | 60 | Sample absent |  |  |
| 13.5 | 3,0,1 | 4 | 40 | 4,2,2 | 10 | 100 | 4,0,1 | 5 | 71 |
| 13.5 | 4,1,1 | 6 | 60 | 4,0,0 | 4 | 40 | 3,0,0 | 3 | 42 |
| 14.5 | 4,1,1 | 6 | 60 | 4,0,0 | 4 | 40 | 4,0,2 | 6 | 85 |
| 15.5 | 4,3,3 | 10 | 100 | 4,2,2 | 8 | 80 | 4,1,2 | 7 | 100 |
| 18.5 | 4,2,2 | 10 | 100 | 4,2,3 | 9 | 90 | Sample absent |  |  |
| **Potential new variants** | | | | | | | | | |
| Embryonic Day | Head | Total intensity | % relative expression | RE | Total intensity | % relative expression | WE | Total intensity |  |
| 12.5 | 4,1 | 5 | 71 | 4,1 | 5 | 71 | Sample absent |  |  |
| 13.5 | 3,1 | 4 | 57 | 4,2 | 6 | 85 | 4,1 | 5 | 83 |
| 13.5 | 4,1 | 5 | 71 | 4,0 | 4 | 57 | 3,0 | 3 | 50 |
| 14.5 | 4,1 | 5 | 71 | 4,0 | 4 | 57 | 4,2 | 6 | 100 |
| 15.5 | 4,3 | 7 | 100 | 4,2 | 6 | 85 | 4,2 | 6 | 100 |
| 18.5 | 4,2 | 6 | 85 | 4,3 | 7 | 100 | Sample absent |  |  |
| **Expected transcript variants** | | | | | | | | | |
| **Embryonic Day** | **Head** | **Total intensity** | **% relative expression** | **RE** | **Total intensity** | **% relative expression** | **WE** | **Total intensity** | **% relative expression** |
| 12.5 | 1 | 1 | 33 | 1 | 1 | 50 | Sample absent |  |  |
| 13.5 | 0 | 0 | 0 | 2 | 2 | 100 | 0 | 0 | 0 |
| 13.5 | 1 | 1 | 33 | 0 | 0 | 0 | 0 | 0 | 0 |
| 14.5 | 1 | 1 | 33 | 0 | 0 | 0 | 0 | 0 | 0 |
| 15.5 | 3 | 3 | 100 | 2 | 2 | 100 | 1 | 1 | 100 |
| 18.5 | 2 | 2 | 66 | 2 | 2 | 100 | Sample absent |  |  |

| ***Phc1*** | | | **Expected band sizes**  **Primer 1- 537, 669, 693bp**  **Primer 2- 500, 507, 389bp**  **Primer 3- 183, 190bp** | | | | **Observed band sizes**  **Primer 1- 669, 537, 300, 200bp**  **Primer 2- 600, 500, 275, 180bp**  **Primer 3- 570 500 190/183bp** | | |
| --- | --- | --- | --- | --- | --- | --- | --- | --- | --- |
| **Cumulative** | | | | | | | | | |
| **Embryonic Day** | **Head** | **Total intensity** | **% relative expression** | **RE** | **Total intensity** | **% relative expression** | **WE** | **Total intensity** | **% relative expression** |
| 12.5 | 7,7,0 | 14 | 82 | 5,3,5 | 13 | 93 | 0,0,0 | 0 | 0 |
| 13.5 | 2,3,4 | 9 | 53 | 6,5,3 | 14 | 100 | 8,3,2 | 13 | 100 |
| 14.5 | 2,10,2 | 14 | 82 | 2,2,4 | 8 | 57 | 2,4,2 | 8 | 62 |
| 15.5 | 5,0,0 | 5 | 29 | 0,0,5 | 5 | 36 | 0,4,0 | 4 | 31 |
| 18.5 | 6,7,4 | 17 | 100 | 4,4,4 | 12 | 86 | 0,0,0 | 0 | 0 |
| **Expected transcript variants** | | | | | | | | | |
| **Embryonic Day** | **Head** | **Total intensity** | **% relative expression** | **RE** | **Total intensity** | **% relative expression** | **WE** | **Total intensity** | **% relative expression** |
| 12.5 | 4,3,0 | 7 | 78 | 0,2,2 | 4 | 80 | 0,0,0 | 0 | 0 |
| 13.5 | 0,1,2 | 3 | 33 | 2,2,1 | 5 | 100 | 0,2,2 | 4 | 100 |
| 14.5 | 1,3,1 | 5 | 56 | 0,1,2 | 3 | 60 | 0,2,0 | 2 | 50 |
| 15.5 | 0,0,4 | 4 | 44 | 0,0,4 | 4 | 80 | 0,3,0 | 3 | 75 |
| 18.5 | 2,3,4 | 9 | 100 | 0,1,2 | 3 | 60 | 0,0,0 | 0 | 0 |

| ***Phc3*** | | | **Expected band size**  **171bp** | | | | **Observed band sizes**  **300, 171bp** | | |
| --- | --- | --- | --- | --- | --- | --- | --- | --- | --- |
| **Cumulative** | | | | | | | | | |
| **Embryonic Day** | **Head** | **Total intensity** | **% relative expression** | **RE** | **Total intensity** | **% relative expression** | **WE** | **Total intensity** | **% relative expression** |
| 12.5 | 4,1 | 5 | 100 | 4 | 4 | 100 | Sample absent |  |  |
| 13.5 | 4,1 | 5 | 100 | 4 | 4 | 100 | 4 | 4 | 100 |
| 13.5 | 3 | 3 | 60 | fail | 0 | 0 | 4 | 4 | 100 |
| 14.5 | 2 | 2 | 40 | 1 | 1 | 25 | 4 | 4 | 100 |
| 15.5 | 4 | 4 | 80 | 4 | 4 | 100 | 4 | 4 | 100 |
| 18.5 | 4 | 4 | 80 | 4 | 4 | 100 | Sample absent |  |  |
| **Expected transcript variants** | | | | | | | | | |
| **Embryonic Day** | **Head** | **Total intensity** | **% relative expression** | **RE** | **Total intensity** | **% relative expression** | **WE** | **Total intensity** | **% relative expression** |
| 12.5 | 4 | 4 | 100 | 4 | 4 | 100 | 0 | 0 | 0 |
| 13.5 | 4 | 4 | 100 | 4 | 4 | 100 | 4 | 4 | 100 |
| 13.5 | 3 | 3 | 75 | 0 | 0 | 0 | 4 | 4 | 100 |
| 14.5 | 2 | 2 | 56 | 1 | 1 | 25 | 3 | 3 | 75 |
| 15.5 | 4 | 4 | 100 | 4 | 4 | 100 | 4 | 4 | 100 |
| 18.5 | 4 | 4 | 100 | 4 | 4 | 100 | 0 | 0 | 0 |

| ***Gapdh*** | **Expected band size**  **248bp** | **Observed band sizes**  **350, 248bp** |
| --- | --- | --- |

**S2 Table- Primer sequences**

**Primers used for PCR amplification**

Primers used for genes were designed by using Ensemble and Primer 3. These designed primers were obtained from the supplier (Macrogen, Korea).

| **Gene ID for primers** | **Primer Sequence** | **Melting temperature (°C)** | **Annealing temperature (°C)** | **Product sizes known** |
| --- | --- | --- | --- | --- |

| Cdk5Rap2_F1  Cdk5Rap2_R1 | GGAGATGACTCTGGCTCTGG  GCTGTCTGGTGACTGCTGAG | 60  60 | 55 | 745bp, 973bp |
| --- | --- | --- | --- | --- |
| Pde4Dip_F1  Pde4Dip_R1 | GCTCAGCAGGGAACTACAGG  GGCTTGGGTATGGAGTGAAA | 60  60 | 55 | 252bp, 405bp |
| Cep63_F1  Cep63_R1 | GAGATGGAGGCTTTGTTGGA  CTGCTTCTCCCAGTCCCAGAG | 60  60 | 56 | 480bp, 302bp |
| Ccdc67_F1  Ccdc67_R1 | ACTGTGCCAGAGTCATGCTG  CAGGTCTTCGTGATTGAGCA | 60  60 | 56 | 423bp |
| Phc1.M.C.2-4F  Phc1.M.C.2-4R | GGCCCCAGATAGCACAGATG  CCTGTCGACTGGCAGCAATC | 60  61 | 55 | 190bp, 183bp |
| Phc1.M.C.2-5.F  Phc1.M.C.2-5.R | GCGAAGGGCTTGAGTCAGA  ATACATCTGGGCCTGGGACT | 59  60 | 54 | 500bp |
| Phc1.M.C.5-7.F  Phc1.M.C.5-7.R | AGTCCCAGGCCCAGATGTAT  GGATCTGTGTGTATGTGGCTGA | 60  61 | 54 | 537bp, 693bp, 667bp |
| Phc3_Q.F  Phc3_Q.R | TTCTCCGCCTTTAACTGTGT  TTTGGCCCTGTTTGAAGAGC | 59  59 | 53 | 171bp |

| **Control primer ID** | **Primer Sequence** | **Melting temperature (°C)** | **Annealing**  **Temperature (°C)** | **Product Length (bp)** |
| --- | --- | --- | --- | --- |
| Gapdh_Q.F  Gapdh_Q.R | GGGTCCCAGCTTAGGTTCAT  CATTCTCGGCCTTGACTGTG | 60  62 | 55 | 248 |
